# From pioneer to repressor: Bimodal foxd3 activity dynamically remodels neural crest regulatory landscape *in vivo*

**DOI:** 10.1101/213611

**Authors:** Daria Gavriouchkina, Ruth M Williams, Martyna Lukoseviciute, Tatiana Hochgreb-Hägele, Upeka Senanayake, Vanessa Chong-Morrison, Supat Thongjuea, Emmanouela Repapi, Adam Mead, Tatjana Sauka-Spengler

## Abstract

The neural crest (NC) is a transient embryonic stem cell population characterised by its multipotency and broad developmental potential. Here, we perform NC-specific transcriptional and epigenomic profiling of *foxd3*-mutant versus wild type cells *in vivo* to define the gene regulatory circuits controlling NC specification. Together with global binding analysis obtained by foxd3 biotin-ChIP and single cell profiles of *foxd3*-expressing premigratory NC, our analysis shows that during early steps of NC formation, foxd3 acts globally as a pioneer factor to prime the onset of genes regulating NC specification and migration by re-arranging the chromatin landscape, opening *cis*-regulatory elements and reshuffling nucleosomes. Strikingly, foxd3 then switches from an activator to its canonical role as a transcriptional repressor. Taken together, these results demonstrate that foxd3 acts bimodally in the neural crest as a switch from permissive to repressive nucleosome/chromatin organisation to maintain stemness and define cell fates.

## Introduction

The forkhead/winged helix transcription factor FoxD3 is an important stem cell factor that functions reiteratively during development. At early stages of development, it is thought to maintain pluripotency of epiblast cells. In embryonic stem cells, its loss leads to premature differentiation into mesendodermal lineages while markers of the ectoderm are reduced (Liu and Labosky, 2008). Later, FoxD3 plays a critical role in the specification and subsequent differentiation of the neural crest (NC), a remarkable transitory and multipotent embryonic cell population. NC cells are specified at the border of the forming central nervous system (neural plate border), but then undergo an epithelial to mesenchymal transition (EMT) to delaminate from the neural tube, migrating into the periphery where they give rise to diverse derivatives such as peripheral ganglia, craniofacial skeleton and pigmentation of the skin (Sauka-Spengler and Bronner-Fraser, 2008). Although individual neural crest cells are multipotent (Baggiolini et al., 2015; Bronner-Fraser and Fraser, 1988), it has been a matter of debate whether the NC population as a whole is homogeneous or a heterogeneous mixture of cells specified toward a particular fate (Harris and Erickson, 2007; Krispin et al., 2010; Nitzan et al., 2013a).

The molecular mechanisms by which FoxD3 influences ES cell development *in vitro* have been extensively studied. During the transition from näive to primed pluripotency cells, FoxD3 represses enhancers by recruiting H3K4 demethylase, Lsd1, resulting in a decrease of active enhancer marks and an increase in inactive enhancer marks (Respuela et al., 2016). During the subsequent pluripotent to epiblast cell transition, FoxD3 primes enhancers by co-recruiting NuRD complex members Brg1 and histone deacetylases 1/2 (HDAC1/2). As a result, different subsets of enhancers get fully activated or are kept repressed during differentiation, depending on the effects mediated by HDAC1/2 removal or retention (Krishnakumar et al., 2016). These studies led to the realisation that FoxD3-mediated gene regulation in ES cells may function via modulation of associated enhancers.

In contrast to ES cells, the molecular mechanisms through which neural crest cells transition from stem cells to fate restricted cells in the embryo and the role of FoxD3 therein remain poorly understood. A neural crest gene regulatory network (NC-GRN) has been assembled that describes the genes expressed during NC ontogeny and their epistatic relationships (Sauka-Spengler and Bronner-Fraser, 2008). Within this framework FoxD3 is known to act downstream of neural plate border (NPB) genes and upstream of factors mediating EMT (Betancur et al., 2010a; Simoes-Costa and Bronner, 2015). In the zebrafish embryo, *foxd3* is one of the earliest zygotically expressed genes (Lee et al., 2013), first detected during epiboly in the dorsal mesendoderm, ectoderm (Wang et al., 2011) and later in the NPB, tailbud mesoderm and floor plate (Odenthal and Nusslein-Volhard, 1998). A second phase of *foxd3* expression occurs in premigratory neural crest cells within the neural folds at all axial levels. Even later, *foxd3* becomes restricted to a subset of migrating cranial neural crest cells and is downregulated in the trunk crest, reappearing in neural crest-derived peripheral glia and other tissues like the somites (Gilmour et al., 2002; Kelsh et al., 2000).

Here, we tackle the molecular mechanisms by which *foxd3* influences neural crest development by taking advantage of wild type and mutant zebrafish *foxd3* lines to characterize the transcriptional and epigenetic landscape of *foxd3*-expressing cells *in vivo*. Intriguingly, we observed a decoupling of the different strategies employed by foxd3 to regulate gene expression over the course of neural crest ontogeny. Inconsistent with its previously defined role as a transcriptional repressor, *foxd3* knockout resulted in global downregulation of neural crest genes. Instead, the results favour the idea that foxd3 acts as a pioneer factor during early stages of neural crest development, as shown by comprehensibly analysing the effects of foxd3 on chromatin accessibility, nucleosome positioning, and transcriptional/epigenetic profiling. At later stages it assumes its canonical role as a transcriptional repressor. We also demonstrate that foxd3 drives several independent chromatin-organising mechanisms, switching from activator to repressor roles to orchestrate multiple regulatory programmes during NC formation, starting with priming early NC specification to fine tuning essential signalling pathways, maintaining stemness by controlling stem cell programmes and preventing untimely migration/differentiation into defined NC derivatives. Moreover, our single cell RNA-seq results show that *foxd3* positive cells display a distinct and homogeneous molecular signature in a stage-specific manner. Our data provide an extensive representation of gene co-expression modules guiding different steps of neural crest development. While respecting the general network architecture of our previously proposed NC-GRN (Sauka-Spengler et al., 2007), this analysis provides exquisite detail of the transitions between modules. By characterising *foxd3*-expressing cells during key stages of NC ontogeny at the single cell, transcriptomic and epigenetic level in both wild type and mutant embryos, our *in vivo* results present the most comprehensive analysis of the NC-GRN to date and provide an integrated platform to interrogate NC gene regulatory circuits.

## Results

### Analysis of foxd3-expressing cells reveals transcriptionally distinct populations

As a first step in characterising the global developmental function of FoxD3, we compared the molecular signatures of *foxd3*-expressing cells (*foxd3*+) to those lacking *foxd3* (*foxd3*-). To this end, we used a genetrap line, *Gt(foxd3-citrine)^ct110^* (Hochgreb-Hägele et al., 2013), which drives expression of foxd3-Citrine fusion, yielding fluorescent signal in endogenous *foxd3*+ cells, enabling their isolation by FACS (Fig. 1A). We evaluated differential gene expression between *foxd3*+ and *foxd3* – populations at three stages of neural crest (NC) development using RNA-seq: (1) prospective progenitors at 75% epiboly; (2) premigratory NC at 5-6 somite stage (ss); and (3) migratory NC at 14-16ss In addition our data were compared with *foxd3*-expressing cells at 50% epiboly, in which neural crest specification had not yet started, using published single cell RNA-seq dataset (Satija et al., 2015) (for details see Fig. S2B).

**Figure 1.**
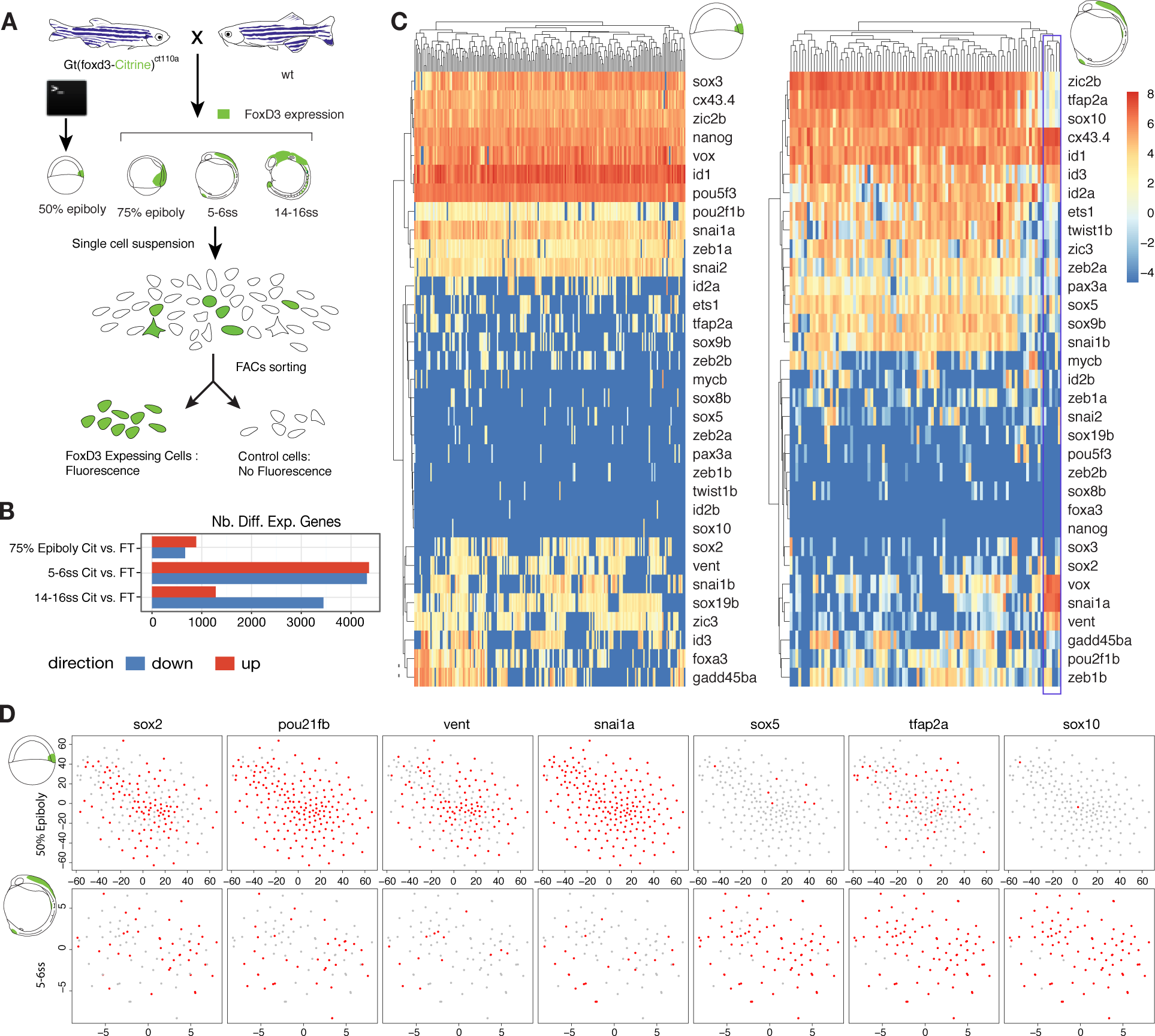
(A) Experimental strategy for obtaining *foxd3*-expressing cells. The genetrap zebrafish line, *Gt(foxd3-citrine)^ct110^*, expressing foxd3-Citrine fusion is outcrossed to wildtype resulting in fluorescent signal in endogenous *foxd3*+ cells, enabling their isolation by FACS. Profiles at 50% epiboly were obtained using a resampling approach similar to bootstrap to generate bulk of *foxd3*+ cells from previously published 50% epiboly single cell (SC) RNA-seq. See Fig. S2. (B) Comparison in number of differentially expressed (enriched and depleted) genes at 75% epiboly, 5-6ss and 14-16ss (FT, flow through). (C) Heatmaps illustrating the hierarchical clustering of *foxd3*+ single cells at 75% epiboly (200 cells) and 5-6ss (93 cells) and showing transcriptional levels of selected NC and stem cell genes. (D) tSNE plots for selected stem cell (*sox2, pou2f1, vent*) and NC genes (*snai1a*, *sox5*, *tfap2a*, *sox10*) indicate individual epiblast and NC cells do not reveal cell subpopulations. Clustering of 5-6ss NC cells singles out a small group of pluripotent non-specified NC progenitors (boxed in purple).

Principal component analysis (PCA) and scatterplots of normalised read counts comparing biological replicates show high level of reproducibility in our experiments (Fig. S1). Differential gene expression was observed at all stages examined, confirming that *foxd3*+ cells constitute a distinct cell population. The highest number of differentially regulated genes was observed at 5-6ss (*p*-value*<*0.05) prior to onset of migration, suggesting that NC transcriptional diversity peaks at premigratory NC stage (Fig. 1B). Starting from 5-6ss *foxd3*+ cells showed significant enrichment of the majority of NC specifiers (*sox9b, sox10, snai1b, twist1b, msxc, pax3a, ednrab* etc.), while 14-16ss *foxd3*+ NC cells were enriched in molecules necessary for active migration (*plexinA3 robo3, tenC, Disc1*) and differentiation into specific NC derivatives such as glia (*olig2, shha/b*), cartilage (*sox9b, acvrIIb-activin receptor IIb, dlx1a, foxl1, dlx2a, fam212aa*) or peripheral neurons (*nkx6, prdm1a, noto, hes12/9, nr4a2b/a, otxb*) (Fig. S2C-F). In contrast, numerous signalling genes (*wnt, fzd, dkk, bmp*, or delta homologues) were depleted, suggesting that, by 14-16ss, actively migrating NC cells repress essential components of major signalling pathways involved in neural crest specification (Fig. S2C). Furthermore, premigratory NC also showed statistically significant enrichment in non-canonical Wnt signalling pathway molecules (*wnt4a, wnt5b, wnt11*) and planar cell polarity (PCP) proteins (*prickle1b, 2b* and *3*), as well as depletion of transcripts coding for vangl PCP proteins 1 and 2, in line with an essential role of non-canonical Wnt/PCP pathway in NC migration in zebrafish and *Xenopus* (De Calisto et al., 2005; Matthews et al., 2008). These analyses confirm specificity of our datasets and further sanction the utility of *foxd3*+ cells for genome-wide studies.

Taken together, differential expression analysis of *foxd3*+ cells reveals major differences between cells at epiboly versus 5-6ss. Whereas the core neural crest specifier genes are actively repressed at epiboly (Fig. S2B,C), the NC specification programme is fully established by 5-6ss, suggesting that it is switched on between these two stages. At later stages (14-16ss), *foxd3*+ cells shut down the signalling pathways involved in specification while activating gene programmes required for cell migration and differentiation (For detailed analysis of the transcriptional landscape of *foxd3*+ cells at all three stages and summary of the results see Fig. S2).

### Single cell RNA-seq reveals homogeneity of foxd3-expressing cells at both 50% epiboly and 5-6ss

There have been debates in the literature regarding whether the premigratory neural crest is a homogeneous or heterogeneous cell population (Harris and Erickson, 2007; Krispin et al., 2010; Nitzan et al., 2013a). To address this *in vivo*, we carried out single-cell RNA-seq (scRNA-seq) at 5-6ss (Fig. S3A). Metrics indicated obtained libraries were of excellent quality (high complexity, a high number of uniquely mapped sequencing reads and up to 5,500 transcripts detected per cell, Fig. S3B). In *Xenopus*, it has been suggested that neural crest cells may retain blastula-stage competence (Buitrago-Delgado et al., 2015). To investigate the transcriptional heterogeneity among single cells, focusing on genes expressed both in premigratory NC and at blastula stages, we found that orthologs of *Xenopus* genes were indeed expressed in the 50% epiboly *foxd3*+ cells in zebrafish (Fig. 1C). We carried out T-Distributed Stochastic Neighbor Embedding (tSNE) clustering of scRNA-seq samples comparing expression of the core NC and stem cell genes in single *foxd3*+ cells at 50% epiboly and 5-6ss (Fig. 1D). At both stages nearly all *foxd3*+ cells expressed the pluripotency factor *cx43.4* and neural plate border specifiers *id1* and *zic2b* at high levels, while more than 50% of cells expressed *pou2f1b*, *zic3* and *id3* (Fig. 1C). Interestingly, however, the expression of core pluripotency factors was different at the two stages examined. The majority of *foxd3*+ single cells at 50% epiboly expressed *Oct4* orthologs (*pou5f3*, *pou2f1b*), *SoxB* ortholog (*sox3*), *nanog* and *vox* (reminiscent of *Xenopus XOct, Xsox2* and *XVent*) (Buitrago-Delgado et al., 2015). In contrast, 5-6ss single *foxd3*+ cells did not express *nanog* and only a few cells expressed *sox3* or *pou5f3* at low levels (Fig. 1C). Furthermore, *foxd3*+ gastrula progenitors expressed a different complement of orthologs of EMT factors compared to premigratory NC, with *zeb1a*, *snai1a* and *snai2* present at 50% epiboly, but poorly expressed in most 5-6ss *foxd3*+ NC cells, which favoured *zeb1b/2a* and *snai1b* (Fig. 1C, D). NC specifiers (*sox5, sox10, twist1b, pax3a*) were expressed at high levels in almost all 5-6ss *foxd3*+ NC cells but were absent from the majority of 50% epiboly *foxd3*+ cells where early NC specifiers (*snai1b, sox9b, tfap2a, ets1, id2a*) were expressed more pervasively (Fig. 1C,D). These findings indicate that while they may use similar regulatory circuits, epiboly-stage and NC-stage *foxd3*+ cells maintain their pluripotency using different sets of paralogous factors.

Further tSNE and PCA clustering of single cell transcriptomes at 5-6ss, based on either all 5,243 or top 500 most divergent genes (Fig. S3C,D) failed to identify multiple specific NC subpopulations, but singled out a small cluster of NC cells which expressed negligent levels of *bona fide* NC specifiers (*zic2b*, *tfap2a*, *sox10*, *twist1b*, *ets1* or *pax3a*) and lower levels of *foxd3* itself. However, these cells expressed high levels of *lratb*, *cxcr4b* and *ved* (Fig. S3E), as well as other markers of stemness (*snai1a*, *vent*, *vox* and *cx43.4*, Fig. 1C,D), suggesting they possibly represent pluripotent non-specified NC progenitors maintained in premigratory NC. Furthermore, clustering and visualising gene expression on selected NC specification gene set (Fig. S3E) points to two NC cell groups with specific hindbrain rhombomere identities (*egr2b*/*hoxa3a*-positive, *hoxb1a*-negative r5 cells and *hoxa2b*/*hoxb1a*-positive, *egr2b*/*hoxa3a*-negative r4 cells) (Trainor and Krumlauf, 2000).

By and large, these results show that both *foxd3*+ epiblast and *foxd3*+ premigratory NC cell populations are non-heterogeneous. Moreover, both populations feature similar pluripotency signature. However, rather than expressing the same genes, premigratory NC cells express paralogous pluripotency factors to those in the epiblast. This suggests that there is redeployment of a paralogous GRN rather than maintenance of the epiblast GRN in the newly specified neural crest. This challenges the proposition that NC cells are residual blastula cells (Buitrago-Delgado et al., 2015).

### Knockout of foxd3 causes genome-wide transcriptional changes

We next inquired how *foxd3* depletion affected NC progenitor cells on a transcriptional level using *Gt(foxd3-mCherry)^ct110^* line, together with the aforementioned *Gt(foxd3-Citrine)^ct110^* line (Fig. 2A,B). In both transgenics, fluorescent reporter proteins interrupt the DNA binding domain, creating mutant fluorescent *foxd3* alleles (Hochgreb-Hägele and Bronner, 2013). These lines were crossed and dissociated embryonic cells obtained from corresponding clutches were FAC-sorted to isolate Citrine only-expressing *foxd3*+ cells(C) as a control, and *foxd3*-mutant cells expressing both fluorophores (Citrine and Cherry, CC) (Fig. 2A,B). We investigated three key stages of neural crest ontogeny (75% epiboly, 5-6ss, and 14-16ss stage) as above (Fig. 1A,B). At 75% epiboly, differential expression analysis between *foxd3*-mutant (CC) and control samples(C) yielded comparable numbers of upregulated and downregulated genes. In contrast at 5-6ss and 14-16ss, a larger number of putatively repressed foxd3 targets (or upregulated genes) was observed (Fig. 2C,D), suggesting a possible change between activator and repressor roles of foxd3 during embryogenesis. Sets of upregulated and downregulated genes were distinct at different stages, with some level of overlap between 5-6ss and 14-16ss, in particular amongst the genes de-repressed upon *foxd3* knockout (Fig. 2E,F).

**Figure 2.**
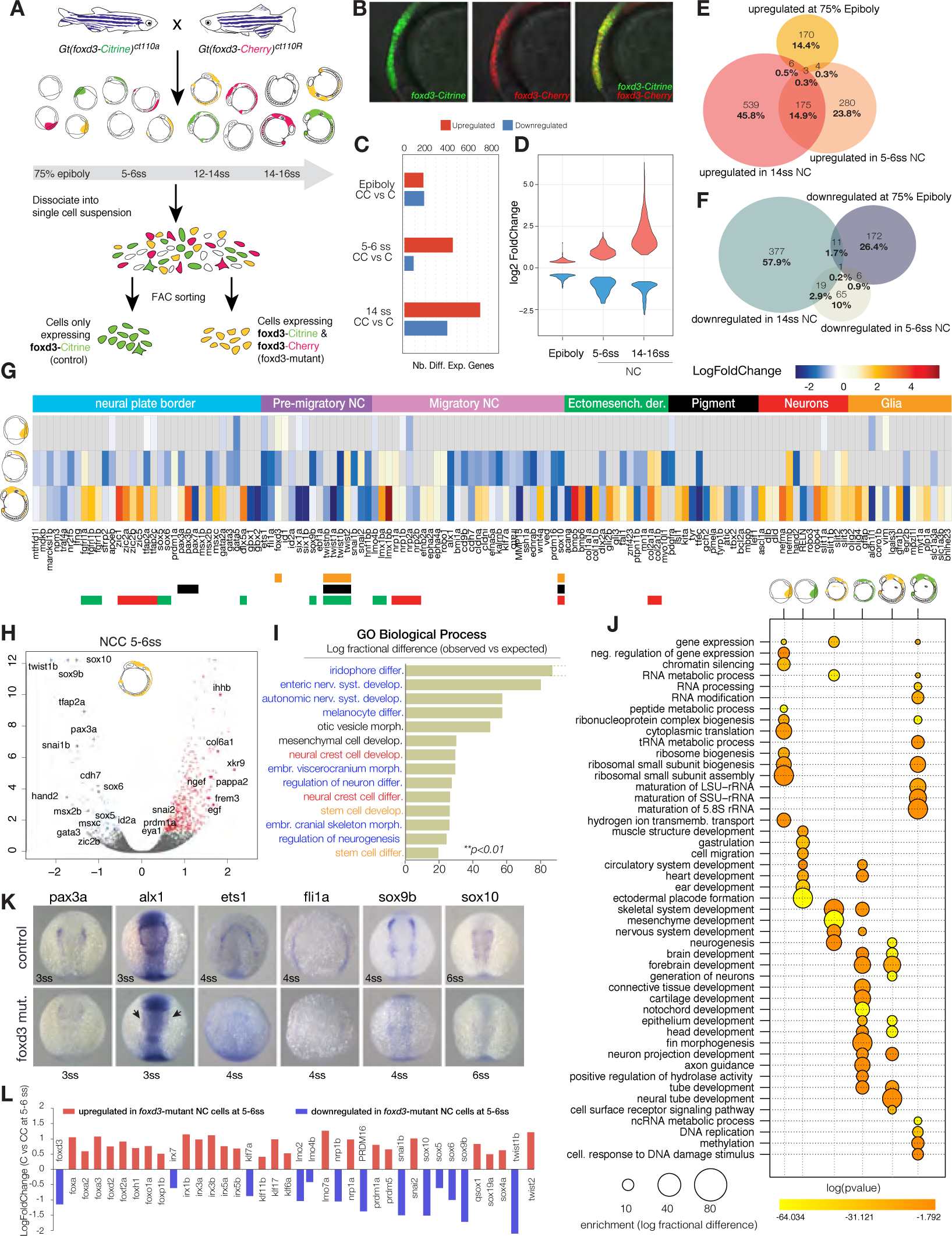
Transcriptional profiling of *foxd3* mutant NC. (A) Experimental strategy for obtaining *foxd3*-mutant (yellow) and *foxd3*-control (green) NC cells. Mutant (Citrine/Cherry, CC) and control (Citrine-only, C) NC cells were isolated by FACS from crosses of heterozygote fluorescent *foxd3* transgenic fish, foxd3-mCherry and foxd3-Citrine at three stages – 75% epiboly, 5-6ss and 14-16ss. (B) Lateral view of a *foxd3*-mutant embryo expressing both Citrine and mCherry instead of *foxd3* in premigratory NC. (C) Bar plot comparing numbers of differentially expressed genes in *foxd3*-mutant and control NC and (D) violin plots comparing fold change differences. (E,F) Venn diagrams comparing upregulated (E) and downregulated (F) genes in *foxd3*-mutant cells. (G) Heatmap showing fold change in expression of known NC genes between *foxd3*-mutant and control cells at 75% epiboly, 5-6ss and 14-16ss. Genes are grouped to reflect NC-GRN structure. (H) Volcano plot highlighting that most NC specifiers are downregulated in *foxd3*-mutant NC at 5-6ss and (I) Bar plot showing gene ontology (GO) terms significantly enriched (\*\**p<*0.01) to downregulated genes, highlighting the roles in stem cell (orange) and NC development (red) and formation of NC derivatives (blue). (J) Bubble plot summarizing enrichment and *p*-values (Benjamini-Hochberg corrected) for the most significant biological process GO terms associated to differentially expressed genes. (K) *In situ* hybridization of 3-6ss zebrafish embryos (dorsal view) showing decrease/loss in expression of NC specifier genes in *foxd3*-mutants. (L) Bar plot representing fold change in expression of NC factors showing that paralogues are differentially regulated by *foxd3*.

FoxD3 has been shown to be required for maintenance of multipotent NC progenitor pool and, at later stages, for control of distinct NC lineage decisions, mostly by repressing mesenchymal and promoting neuronal derivatives (Dottori et al., 2001; Kos et al., 2001; Lister et al., 2006; Montero-Balaguer et al., 2006; Mundell and Labosky, 2011; Stewart et al., 2006; Teng et al., 2008). Examination of GO terms over-represented in differentially regulated genes indicated that at 75% epiboly, foxd3 promoted cellular specification by repressing epigenetic mechanisms that silence/compact chromatin, as well as negative regulators of maintained pluripotency (e.g. H2A variants, *id1*, *her3*, *p*-value*<*0.01) (Fig. 2J), whilst at the same time activating tissue-specific programmes, as well as genes controlling progenitor adhesion and migration (e.g. *nrp2a, nrp1b, slit1a, cdh6, itga5*, *p*-value*<*0.05, Fig. 2G,J). Strikingly at 5-6ss, *foxd3*-mutant cells (CC) downregulated a large proportion of known NC genes distributed across all defined NC-GRN modules (*p*-value*<*0.01, Fig. 2G), a large proportion (*~*40%) of which were *bona fide* NC transcription factors (Fig. 2H). Statistical overrepresentation of the downregulated genes yielded significant association with neural crest and stem cell development GO terms (*p*-value*<*0.01, Fig. 2I) and included a multitude of NC derivative fates (Fig. 2I,J) suggesting that foxd3 controlled NC specification events at premigratory stages. Some factors previously reported to act upstream of *foxd3*, such as *prdm1* and *tfap2a/c*, (Fig. 2G) (Li and Cornell, 2007; Powell et al., 2013; Sauka-Spengler and Bronner-Fraser, 2008) were downregulated, challenging the proposed epistatic relationships within the NC-GRN. Downregulation of NC specifiers was confirmed by *in situ* hybridisation (Fig. 2K).

Analysis of *foxd3* mutant cells (CC) at migratory NC stages, showed dysregulation of neural plate border and derivative markers (Fig. 2G,J). Surprisingly, by 14ss we observed an untimely upregulation of markers of ectomesenchymal derivatives (*lmx1ba/b, bmp5/6, col2a1a/b*), and neuronal lineages (*delta b/d, robo4, slit1a/b, slit2/3*), but only partially of melanophore, xanthophore (*isl1, kita1, pmela/b, tyrp1b, ascl1a*) and glial lineages (*gfap. olig2/4, gfra1b, myt1a/b, plp1, slc1a3b, bhlhe23*), which normally would be expressed much later or not expressed in *foxd3*+ NC derivatives (Fig. 2G). Several derivative and ectomesenchymal markers (*col2a1a/b, lmx1bb/b*) and cell surface signalling molecules (*epha4a, slit2/3, robo1/4*), were already de-repressed at premigratory stage (Fig. 2G), in line with the role of foxd3 in preventing premature differentiation into NC derivatives. Statistical over-representation tests associated the upregulated genesets at both 5-6ss and 14ss show multiple GO terms reflecting either dysregulation of derivative formation (neurogenesis, axonogenesis, etc.), or biological processes essential for NC migration (cell surface signalling pathways, cell-cell adhesion) (Fig. 2J). It is of note that derivative markers dysregulated in *foxd3*-mutant cells were not expressed above background in *foxd3*+ control cells (FPKM*>*1) at any stage examined.

Interestingly, several paralogs belonging to same gene family were differentially regulated. For instance, key NC factors (*snai1b*, *twist1b*, etc.) were downregulated in mutant cells, whilst *snai2* and *twist2* were up-regulated (Fig. 2L) offering a possible mechanism for rescue of *foxd3* transcriptional phenotype by paralogous genes. Additionally, several *Fox* transcription factors were upregulated in *foxd3* mutants, which suggests a different, upstream compensating mechanism by different fox genes. Under some conditions, FoxD3 has been shown to auto-repress itself (Hromas et al., 1999; Lister et al., 2006; Pohl and Knochel, 2001). In mutant cells, *foxd3* was upregulated at 75% epiboly, downregulated at 5-6ss and subsequently upregulated at 14-16ss, indicating differential feedback loops controlling *foxd3* expression at different stages of development.

Altogether, these results show that foxd3 plays different regulatory roles depending on the time and context. Importantly, in the absence of a functional foxd3 protein, much of the NC specification module is absent at 5-6ss. While genes associated with NC and stem cell processes are downregulated in the mutant premigratory NC, genes governing migration and differentiation are upregulated at migratory stages, suggesting that foxd3 switches from an activator to a repressor. We also find unexpected compensation in the mutant by other Fox proteins and alternative NC genes that partially rescue NC specification.

### foxd3 affects chromatin accessibility and identifies distal cis-regulatory elements

The counter-intuitive finding that a large number of NC specification factors were downregulated in *foxd3*-mutant at 5-6ss raises the intriguing possibility that, much like FoxA1/A2 factors during endodermal specification (Iwafuchi-Doi et al., 2016), FoxD3 may act as a pioneer factor during NC specification. In this scenario, FoxD3 may modulate the local epigenetic state of multiple *cis*-regulatory elements and thus positively control many NC genes in addition to its canonical role as a transcriptional repressor (Xu et al., 2007; Xu et al., 2009). To assess chromatin accessibility status in *foxd3*-mutants, we carried out cell-type specific ATAC-seq at different stages of NC formation on either FAC-sorted *foxd3*-expressing (C) and *foxd3*-mutant NC cells (CC) (75% epiboly and 5-6ss) or 1-2ss anterior embryonic cells dissected from *foxd3*-mutant and control (CC and C) embryos. In addition, we used our previously published 16ss *sox10*-specific ATAC-seq (Trinh et al., 2017), containing an extensive cohort of *cis*-regulatory elements open in migratory NC.

We recovered a constant number of open chromatin regions (ATAC-seq peaks) at all stages, and their distribution as distal (intronic, intergenic) or proximal (promoter) remains similar. However, the total number of open elements dramatically increased in late migratory and differentiating NC cells, this increase was entirely accounted for by novel distally located elements (Fig. 3A). The knockout of *foxd3* did not affect the distribution of peaks according to genomic annotation (*p*-value = 0.8743 and 0.614 for epiboly and 5-6ss, respectively). Over 60% of total peaks observed in *sox10*-specific differentiated cells were already opened at earlier stages (Fig. 3B,E). To verify whether the chromatin state of promoters and distal *cis*-regulatory elements correlates with gene expression, we compared expression levels of the closest genes to open chromatin regions. We noted a bimodal distribution of gene expression levels associated with putative enhancer elements at all stages, but with putative promoters only at epiboly. Unimodal distributions after epiboly for genes associated with putative promoters indicates a *cis*-regulatory role for foxd3 begins by 5-6ss (Fig. 3C). Moreover, while at 75% epiboly the difference in number of unique peaks in control (C) and mutant (CC) cells is negligible (21% vs 19%), by 5-6ss, the number of peaks in control cells is almost two-fold of that in mutants (33% vs 17%) (Fig. 3D).

**Figure 3.**
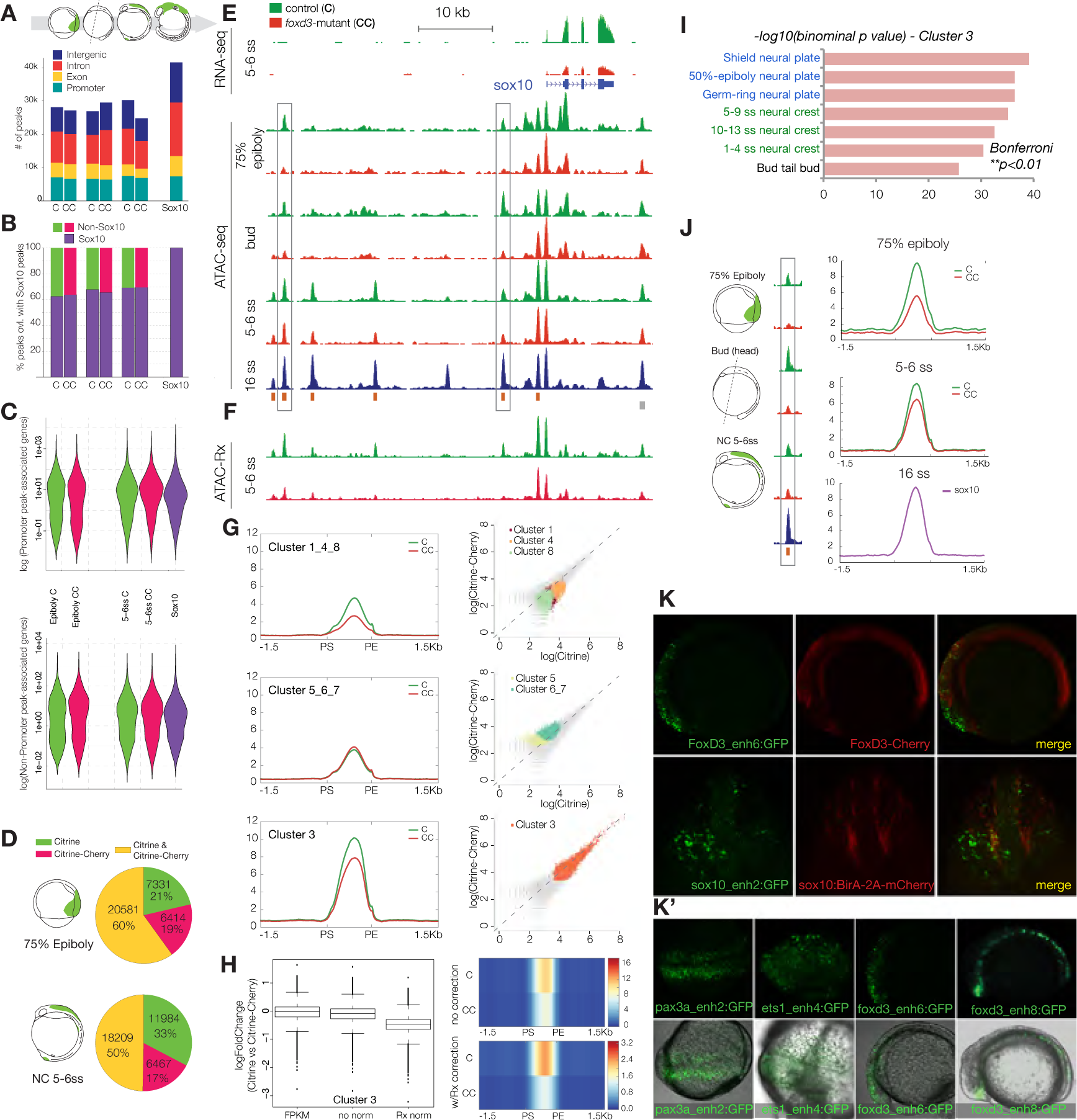
Epigenomic profiling of chromatin accessibility in *foxd3* mutant NC across developmental time. (A) Stacked bar plots depicting genomic annotation of ATAC-seq peaks across stages analysed (75% epiboly, bud stage, 5-6ss and 14-16ss) and (B) quantification of open elements at earlier stages as a proportion of accessible elements detected in migrating/differentiating NC.(C) Violin plots correlating putative promoter and *cis*-regulatory elements with gene expression levels. Bimodal distribution of gene expression is associated with putative enhancers at all stages, but with promoters only at epiboly. (D) Pie charts comparing Citrine-only, Cherry-only and Citrine/Cherry peak number proportions of ATAC-peaks. (E) Genome browser screenshot showing RNA-seq and ATAC-seq profiles in *foxd3* mutant (red) and *foxd3*-control cells (green) within *sox10* locus. (F) Tracks showing normalized ATAC-Rx profiles obtained using reference exogenous *Drosophila* epigenome. (G) Mean density maps of merged profiles and corresponding scatterplots of raw counts for *k*-means clusters featuring elements with differential accessibility and signal levels in *foxd3*-mutant and controls at 5-6ss (H) Box plots and heatmap showing fold change in accessibility and comparing ATAC signal levels between control (C) and mutants (CC) Cluster 3 elements with and without Rx normalisation. (I) Bar chart depicting functional annotation of *k*-cluster 3 shows enrichment in zebrafish gene expression ontology terms linked to NC and neural plate development (Bonferroni \*\**p<*0.01). For further analysis of *k*-cluster see Fig. S4. (J) Merged profiles for 3,565 elements open at 75% epiboly showed more prominent accessibility defect than at 5-6ss (C *>>*CC, *>*50%), suggesting biological compensation over time. (K,K′) *Cis*-regulatory elements from Cluster 3 show NC-specific reporter activity. (K) Lateral and frontal view of embryos injected with *foxd3*-enh6 and *sox10*-enh2 GFP reporter constructs into the genetic background of foxd3-Cherry and sox10:BirA-2A-Cherry, respectively. (K′) Fluorescent and brightfield overlay images of *pax3a* and *ets1* (dorsal view) and *foxd3* (lateral view) enhancers.

To investigate whether dynamics of opening of distal regulatory elements could account for the drastic depletion of NC specification genes at 5-6ss, we compared the ATAC-seq profiles in *foxd3*-mutant (CC) and *foxd3*-expressing control cells (C) (Fig. 3E). *K*-means clustering identified 8 cohesive groups of elements observing three general trends: (1) Clusters 1-4-8 contained lower signal elements with prominent accessibility differences between mutant and controls (C*>>*CC) (2) Clusters 5-6/7 comprised elements of equally low comparable accessibility (C*~*CC) and (3) Cluster 3 contained highly accessible regions with broad ATAC-seq peak distribution that showed intermediate signal decrease in mutant (C*>*CC) (Fig. 3G). Functional annotation of *k*-clusters using GREAT Tool (McLean et al., 2010) singled-out two clusters reflecting NC regulatory mechanisms – Clusters 3 and 4 showed specific enrichment of zebrafish gene expression ontology terms linked to NC and neural plate development (Bonferroni \*\**p<*0.01, Fig. 3I, S4A).

To quantify the observed difference in ATAC-seq signal we adapted a ChIP-Rx method (Orlando et al., 2014) that enables genome-wide quantitative comparative analysis of histone modification ChIP signal (ATAC-Rx). To this end, ATAC was performed on mutant (CC) and control (C) *foxd3*-expressing NC cells at 5-6ss, spiked with *Drosophila melanogaster* S2 cells as a reference exogenous epigenome (Fig. 3F). Quantification after Rx normalisation demonstrated a discernible fold-change difference in accessibility between control (C) and mutant (CC) elements (Fig. 3H), thus further confirming the defect in opening of specific distal *cis*-regulatory elements in *foxd3*-mutant, previously identified by *k*-means clustering.

To investigate dynamics of chromatin opening over developmental time, we performed *k*-means clustering of the 75% epiboly and bud stage ATAC data. We found a subset of Cluster 3 elements was open at 75% epiboly (*~*20%, 3,565 el.), with a more prominent change in enhancer accessibility in *foxd3* mutants at this stage (C*>>*CC, *>*50%) as compared to 5-6ss (Fig. 3J), suggesting the defect in *foxd3* mutants is compensated over time. Using an efficient reporter assay in zebrafish, we tested the activity of 30 putative regulatory elements from Clusters 3 and 4. Cluster 4 regions were not active at 5-6ss, but were possibly used at later stages, to maintain NC specifiers that remained downregulated in 14ss *foxd3* mutants. Cluster 3 elements drove reporter expression at 5-6ss with striking NC-specific activity, recapitulating endogenous expression of their cognate genes (Fig. 3K, K′), thus strongly suggesting they act as their *cis*-regulatory elements.

### Hotspot enhancers associated with downregulated NC specification genes harbour specific NC regulatory code

Cluster 3 included elements involved in both neural and NC development (Fig. 3I). However, *foxd3*-mutants presented defects only in NC formation, suggesting that neural *cis*-regulatory modules may not require foxd3 activity for proper function. Further *k*-means clustering of Cluster 3 revealed two pooled subclusters (Fig. 4A-C): (1) Cluster 3.1 containing of *cis*-regulatory elements that displayed lower accessibility and (2) Cluster 3.2 containing elements with no change in accessibility in *foxd3* mutants. Clusters 3.1 and 3.2 were generated by assembling Cluster 3 subclusters that exhibited similar accessibility characteristics. GREAT analysis further functionally segregated these subclusters: Cluster 3.2 was associated with ontology terms linked only to neural plate/tube development while Cluster 3.1 contained hotspot enhancers implicated in NC specification or neuronal differentiation (Bonferroni, \*\**p<*0.01, Fig. 4D,F).

**Figure 4.**
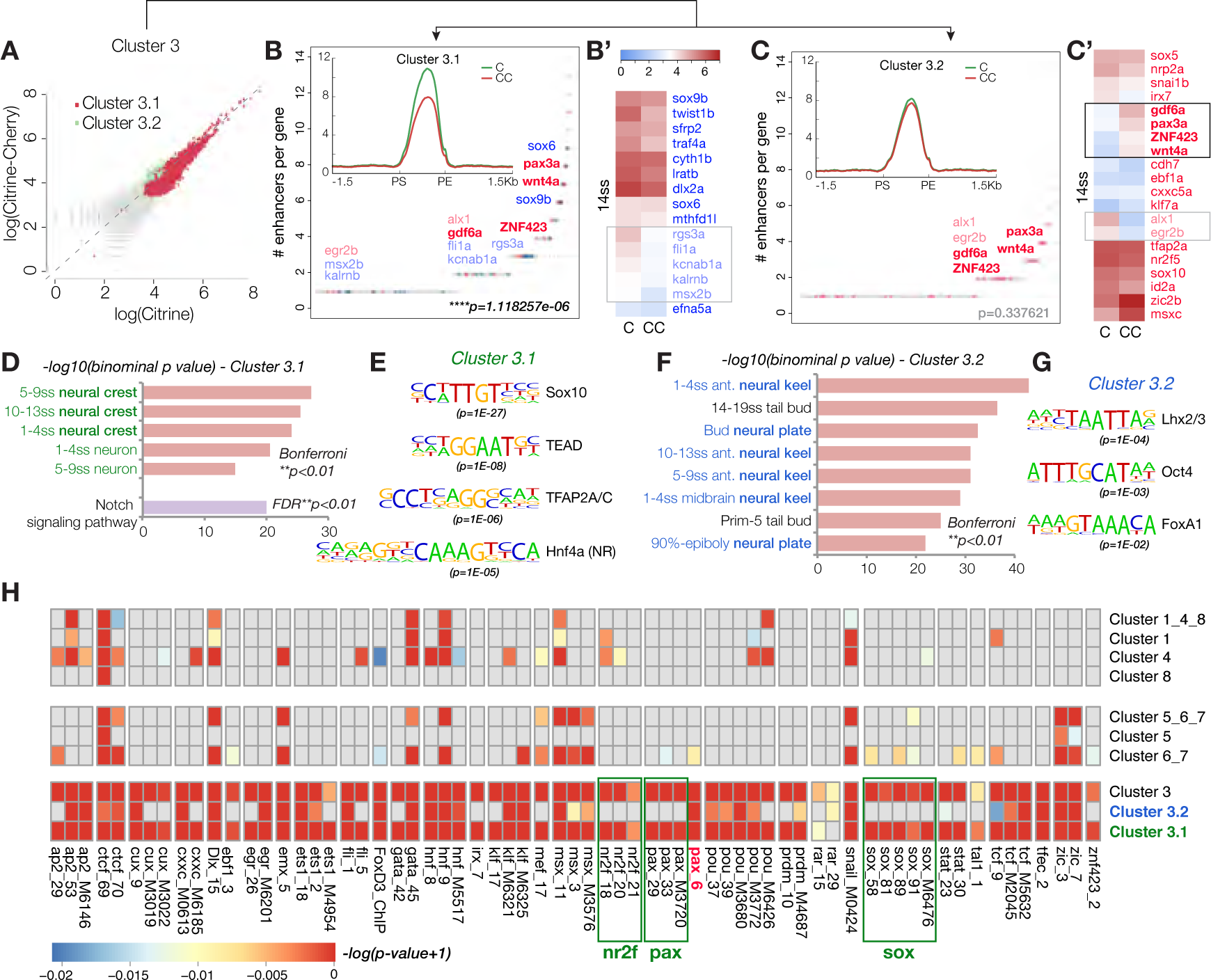
Characterisation of hotspot enhancers. (A) Scatterplot showing subclustering of Cluster 3, one containing elements of lower accessibility in *foxd3* mutants (Cluster 3.1, 12,366 el., R_Cl3.1_=0.77) (B) and the other elements with no change in chromatin accessibility (Cluster 3.2, 4,754 el., R_Cl3.2_=0.97) (C). (B, C) Plots representing genes assigned to Cluster 3.1 (B) and Cluster 3.2 (C) ranked by the number of associated elements. (B′,C′) Heat maps showing later expression (14ss) of NC genes depleted in 5-6ss mutant NC. Genes controlled solely by 3.1 elements (in blue) are shown in (B′) and those harbouring both 3.1 and 3.2 elements (in red) are depicted in (C′). Genes that remain downregulated at 14ss are labelled in light colour print and those overexpressed are shown in bold. (D, F) Functional annotation by GREAT associates Cluster 3.1 with neural crest specification or neuronal differentiation (D) and Cluster 3.2 with neural plate/tube development (F) (Bonferroni \*\**p<*0.01). (E, G) Top Transcription Factor Binding Site (TFBS) motifs enriched in 3.1 (E) and 3.2 (G) elements. (H) Comprehensive TF binding motif map representing significantly enriched TFBS for TF expressed at 5-6ss across different *k*-clusters.

To link the putative regulatory elements, identified in *foxd3*-mutant (CC) and control (C) NC cells to their transcriptional programmes, we first assigned all identified non-promoter ATAC-seq elements to the genes expressed at each corresponding stage (Fig. S4C). To connect the transcriptional and regulatory *foxd3* phenotypes at 5-6ss, we assigned putative enhancers from Clusters 3.1 and 3.2 to the corresponding genes expressed at this stage and ranked those genes by the number of elements associated (Fig. 4B,C). The elements from Cluster 3.1 correlated to the ensemble of NC specification genes downregulated at 5-6ss with high statistical significance (\*\*\*\**p*=1.12E-60). Moreover, no other *k*-cluster, including 3.2, showed significant association to genes either up- or downregulated in *foxd3*-mutant at 5-6ss.

Many NC specifiers that were downregulated in *foxd3*-mutants at 5-6ss recovered their expression by 14ss. We inquired whether Cluster 3.2 regulatory elements (unaffected by loss of *foxd3*) could act instead of hotspot Cluster 3.1 enhancers to rescue cognate gene expression. We found that genes controlled by both Cluster 3.1 and 3.2 elements (Fig. 4C′) compared to those solely using Cluster 3.1 elements (Fig. 4B′) did not recover more efficiently (50% versus 40% of genes, respectively, remained downregulated at 14ss). Instead, an important fraction (*~*25%) of downregulated NC specifiers harbouring 3.2 elements, were, in fact, upregulated by 14ss and such upregulation was not observed for genes solely controlled by Cluster 3.1 hotspot activating enhancers. Moreover, genes differentially upregulated at 14ss associated to Cluster 3.2 with high statistical significance (*p*=4.73E-74), suggesting that 3.2 elements, enriched in FoxA binding motifs (Fig. 4G), were in fact linked to *foxd3*-mediated repression. In contrast, those genes regulated by multiple hotspot 3.1 enhancers (such as *sox6* and *sox9b*) were rescued more efficiently, suggesting an additive effect.

In line with their predicted assigned functions, Transcription Factor Binding Site (TFBS) analysis using Homer suite (Heinz et al., 2010) revealed that Clusters 3.1 and 3.2 harboured distinct regulatory codes. Cluster 3.1 enhancers presented a canonical neural crest signature featuring *bona fide* NC master regulators Sox10 (Sauka-Spengler and Bronner-Fraser, 2008), TFAP2a and nuclear receptor NR2 (Rada-Iglesias et al., 2012) as top enriched binding motifs (Fig. 4E), while Cluster 3.2 top enriched motifs were Lhx2/3, TF involved in neural development and cortical neurogenesis (Bery et al., 2016), Oct4-Sox2 and multiple FoxA motifs (Fig. 4G). Interestingly, the only other *k*-clusters that were enriched in NC motifs (TFAP2a and Ets1, but not Sox10) were clusters 1-4-8 (Fig. S4), suggesting that regulatory elements whose opening is dependent on foxd3 display unifying features of a NC enhancer.

Given that there is a paucity of available zebrafish TFBS, we formulated a novel clustering method to build a comprehensive TF binding motif maps for each enhancer cluster to be used in statistical enrichment analyses. The heatmap of representative enriched DNA motifs showed that the majority of NC TF motifs are present in the hotspot 3.1 neural cres enhancers (Fig. 4H). Cluster 3.2 elements clearly lacked sox10, nr2f and most pax motifs, except for a single pax cluster, comprising of human TF binding motifs for Pax3 and Pax7, previously shown to control both NC, neuronal and mesenchymal derivatives (Manderfield et al., 2014; Murdoch et al., 2012). Moreover, 3.2 enhancers harboured the majority of hnf, tcf, klf, zic and pou motifs, suggesting these elements could both drive NC derivative as well as stem cell maintenance programmes at later stages of NC development.

This analysis singled out 3.1 as the *bona fide* NC enhancer cluster that contained hotspot *cis*-regulatory modules driving NC specification genes at premigratory stages. Defects in the chromatin accessibility of these hotspot enhancers resulted in the decrease of NC specifiers expression in *foxd3* mutants.

### foxd3 primes late regulatory elements used in migratory NC

To quantitatively evaluate events of chromatin opening at 5-6ss, we performed differential accessibility analysis using DiffBind package (Stark R, 2011). We identified 900 peaks that were differentially upregulated in *foxd3*-control (C) versus *foxd3*-mutant (CC) neural crest (Fig. 5A); these elements exhibited low signal at 75% epiboly, only started to open at 5-6ss, but were clearly accessible in the NC at 16ss (Fig. 5B-D). Functional annotation of identified elements by GREAT revealed significant enrichment of GO terms for stem cell development/differentiation, neural crest differentiation/migration and mesenchymal cell differentiation (\*\**p<*0.01) as well as gliogenesiis (\**p<*0.05), further suggesting these regions may act as *cis*-regulatory elements at later stages of NC ontogeny (Fig. 5E). Interestingly, assigned genes included cell adhesion and migration factors that were de-repressed in *foxd3* mutant NC at later stages. Conversely, other associated NC regulatory factors that drive specific NC lineages and are normally highly expressed at later stages were depleted in *foxd3* mutant at 14ss (Fig. 5A). These results clearly suggest that, in addition to NC specification programme at premigratory NC stages, foxd3 continues to aid opening of the *cis*-regulatory elements associated with NC differentiation, while, at the same time, negatively controlling gene expression of cell surface and migration machinery, that ultimately has to be deactivated in order for cells to settle and differentiate. Importantly, association of stem cell development/differentiation genes to late NC enhancers further supports a role for foxd3 in controlling stem cell identity in the migrating/differentiating NC.

**Figure 5.**
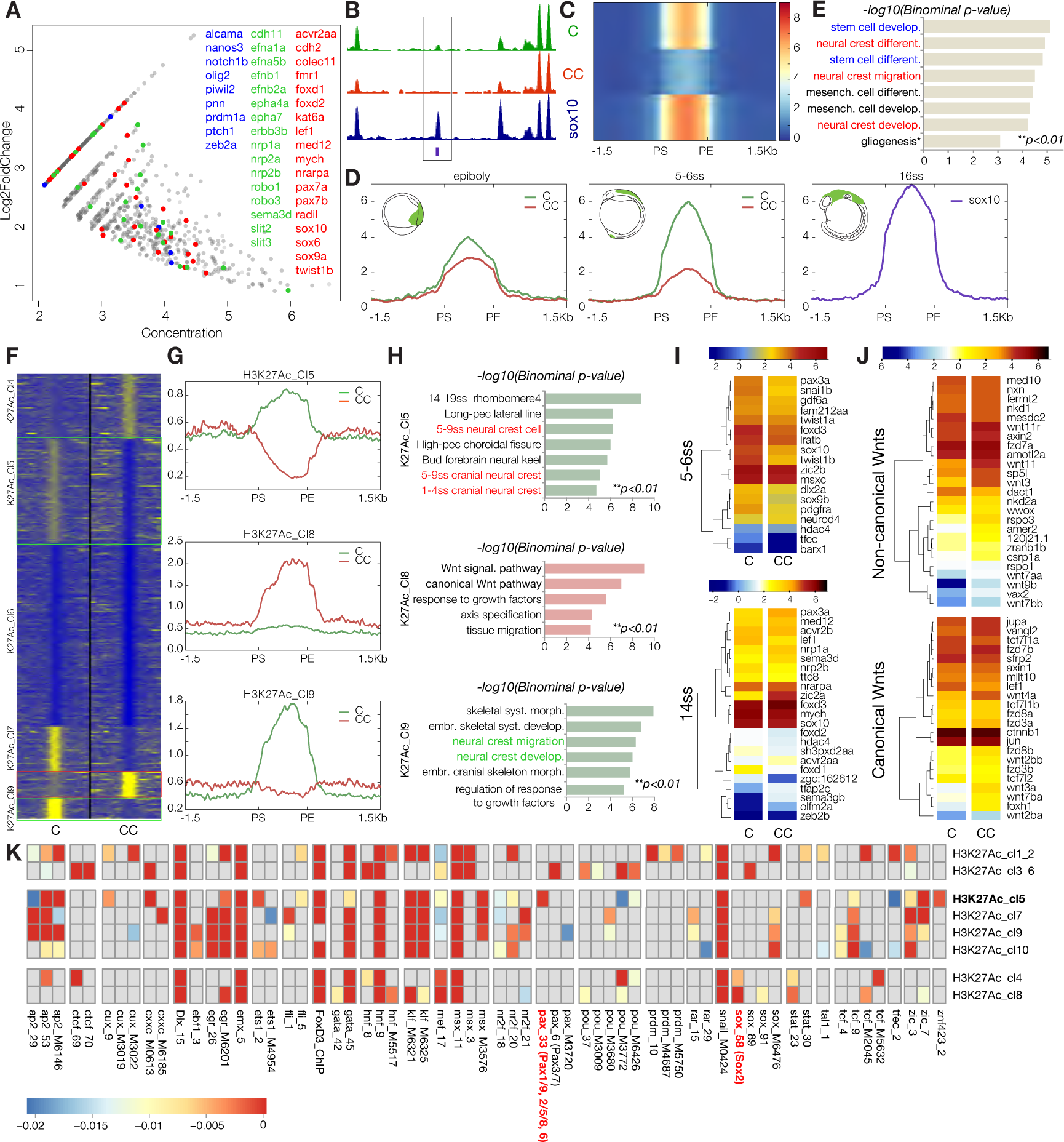
Differential ATAC-seq analysis and clustering of enhancers based of H3K27Ac profiles. (A) Annotated MA plot depicting late opening enhancers significant by Diffbind analysis (*p*-value*<*0.05, FDR*<*0.1) of the ATAC-seq signal at 5-6ss with annotated associated genes (stem cell genes, blue; cell adhesion/migration cues, green; NC specification/differentiation, red).(B) Genome browser screenshot exemplifying type of element isolated by Diffbind (boxed). (C) Heatmap of all elements and (D) collapsed merged profiles indicating that identified elements are closed at epiboly and only start to open at 5-6ss. (E) Functional annotation of Diffbind-identified enhancers shows association with later roles in NC (\*\**p<*0.01). (F) Heatmap depicting *k*-means linear enrichment clustering of H3K27Ac signal across non-promoter ATAC-seq peaks in *foxd3*-mutant (CC, Citrine/Cherry) and control (C, Citrine) at 5-6ss, (G) associated mean merged profiles for selected clusters and (H) corresponding ontology enrichment bar-plots indicating functional role of selected clusters. (I) Heatmaps showing expression of NC specification genes associated with K27Ac Cl5 at 5-6ss and NC migration/differentiation genes associated with K27Ac Cl9 at 14ss in *foxd3*-mutant (CC) and control cells (C). (J) Heatmap depicting expression at 14ss of canonical and non-canonical Wnt pathway molecules associated with K27Ac Cl8 that displays an increase in enhancer K27 acetylation in mutants. (K) TF binding motif map representing significantly enriched TFBS for TF expressed at 5-6ss across different K27Ac-clusters. See Fig. S5 for other K27Ac clusters.

Collectively, our findings demonstrate that foxd3 controls NC gene activation by acting at a *cis*-regulatory level both during early NC specification and at later migratory NC stages. This realisation contributes to converging evidence that foxd3 plays multiple, sometimes opposing roles, particularly during the transition from NC specification to migration/differentiation stages.

### H3K27Ac on NC enhancers is altered in foxd3 mutants

To examine whether H3K27 acetylation, a hallmark of active enhancers, was affected in *foxd3*-mutant NC at 5-6ss, we carried out H3K27Ac ChIP using FAC-sorted *foxd3*-mutant (CC) and control *foxd3*-expressing NC cells (C). *K*-means clustering of H3K27Ac signal identified 10 clusters with patterns of differential H3K27 acetylation on putative *cis*-regulatory elements (Fig. 5F,G; S5). Four clusters (K27Ac Clusters 1, 2, 3 and 6) contained elements with no differential H3K27 acetylation, whereas four clusters showed a decrease (K27Ac Cl5, 7, 9 and 10) and two an increase (K27Ac Cl4 and Cl8) in H3K27Ac signal in *foxd3* mutant NC. In K27Ac Cl5 acetylation in *foxd3*-mutants was abrogated below background levels, possibly indicating active removal of the H3K27Ac mark from the enhancers when they are not primed or bound by NC specific TFs. Functional annotation of this cluster yielded specific enrichment of zebrafish expression GO terms linked to early (premigratory) NC, as well as nervous system development (Bonferroni \*\**p<*0.01, Fig. 5H). The majority of NC genes downregulated in *foxd3* mutants at 5-6ss were associated with one or more K27AC Cl5 elements (*p*=1.59E-05, Fig. 5I), suggesting that some of the enhancers initially opened by foxd3 and used during early NC specification also depended on this factor for appropriate acetylation. Interestingly, K27Ac Cl5 elements also associated with high statistical significance to the genes upregulated both at 5-6ss and at 14ss (*p*=2.87E-05 and *p*=1.19E-05, respectively), mainly involved in the central nervous system development and neuron axonogenesis (Fig. 2I). Similarly, H3K27Ac Cl9 elements, characterised by strong K27Ac signal in controls and defect in *foxd3*-mutant cells (Fig. 5G), were mainly associated with factors regulating late NC events like migration and differentiation into derivatives such as cranial skeletal elements (Fig. 5H). Interestingly, a number of these genes were upregulated in *foxd3*-mutants by 14ss, indicating a supplementary *foxd3*-linked gene/enhancer regulatory mechanism (Fig. 5I). The putative role of foxd3 in repression of these NC genes until post-migratory stages is reminiscent of the observations made in studies of Foxd3 function in germ and pluripotent stem cells (Krishnakumar et al., 2016; Respuela et al., 2016).

In contrast, the increased acetylation in *foxd3*-mutants on elements from cluster K27Ac Cl8 (Fig. 5G) indicated a possible involvement of foxd3 in active repression of this histone modification. GREAT analysis suggested a functional role for these elements in the regulatory control of Wnt signalling pathway components. Both canonical Wnt signalling ligands (*Wnt1,3,3a,8a/b, 10a/b*), receptors (*fzd3,8b,10,fzdb,sfrp1a*), signal transduction effectors (*apc, axin2, wntless, tcf3a/b, tcf15*) as well as non-canonical Wnt signalling ligands (*wnt4a,5b, 7b,11,11r,16*) and signal transduction effectors (*daam1a/b, rho, plc,nfat3b*), while not necessarily differentially expressed in *foxd3*-mutants at 5-6ss, were upregulated at 14ss (Fig. 5J).

Interestingly, DNA motifs enrichment patterns identified in individual K27Ac clusters differed from binding maps of hotspot 3.1 enhancers (Fig. 5K). For instance, enhancers from K27Ac Cl5 that show complete depression in H3K27Ac signal in the absence of *foxd3* are not enriched in sox, prdm or pax3/7 motifs, but contain motifs for other paxs (such as pax2/5/8), and retain ets binding motifs. In addition, similar to other clusters whose elements are acetylated in a *foxd3*-dependent manner (K27Ac Cl7, 9 and 10), K27AC Cl5 enhancers are enriched in tfap2, nr2f and zic motifs. Regions of low acetylation across ATAC peaks in K27Ac Cl3 6 are enriched in CTCF binding motifs, while elements from clusters K27Ac Cl4&8, which may normally require foxd3 binding for maintenance of repressive state, are enriched in binding motifs for neural/stem cell gene sox2.

These results show that foxd3′s effects on H3K27 acetylation of enhancers are context dependent. While correlating positively with H3K27Ac deposition on enhancers of early specification and late fate commitment genes, foxd3-dependent active H3K27Ac removal is associated with repression of Wnt signalling genes.

### Biotin-ChIP confirms direct action of foxd3

To interrogate the sites of direct foxd3 binding at the time of NC specification, we used our recently developed binary biotagging approach (Trinh et al., 2017), enabling specific biotinylation of target proteins of interest *in vivo* for subsequent use in biochemical procedures (Fig. 6A). To this end, the effector transgenic zebrafish line, *TgBAC(foxd3-Avi-2A-Citrine)^ox161^*, expressing Avi-tagged foxd3 protein in endogenous fashion (Fig. 6B), was crossed to the ubiquitous BirA driver, *Tg(ubiq:NLS-BirA-2A-Cherry)^ox114^*, expressing biotin ligase BirA targeted to the nucleus (Fig. 6A) and resulting embryos were collected for biotinChIP-seq (Fig. 6C). BirA-only expressing embryos were used as a control. BiotinChIP-seq revealed 9231 foxd3-bound positive regions in 5-6ss NC cells expressing biotinylated foxd3 and only 10 peaks for the BirA-only control, none of which were associated with NC genes. Approximately 60% of foxd3-bound regions were located within the non-coding regions (intergenic and introns) with 30% peaks overlapping promoters (Fig. 6D). Interestingly, we found that 40% of foxd3-bound regions overlapped with the accessible elements identified using ATAC-seq in *foxd3*-expressing (Citrine-positive NC cells). Of those, a majority (60%) belonged to *k*-means Cluster 3, containing hotspot enhancers that controlled expression of NC specifiers at 5-6ss, whereas another 17% and 20% of foxd3-bound sites were found in the Clusters 1 4 8 and 5 6 7, respectively. Analysis of foxd3-bound regions using MEME suite (Bailey et al., 2015), followed by searches for statistically significantly similar motifs in the known databases using TOMTOM tool (Gupta et al., 2007) identified differences in top position weight matrices (PWMs) and associated binding profiles between the foxd3-bound elements that overlap putative enhancer elements (ATAC-seq peaks) and those falling into regions of inaccessible chromatin (Fig. 6E). *Bona fide* fox motifs were predominantly featured within the ChIP-ed open regulatory regions, and in some cases were combined with partially overlapping consensus binding sites for core neural crest (Sox) and stem cell TFs (HD/Pou) (Fig. 6D), whereas the top foxd3-bound motifs within the closed chromatin were those bound by MADS-box TF MEF2 and hormone-nuclear receptor NR2/3, in addition to consensus motifs for different fox factors including foxds (Fig. 6E).

**Figure 6.**
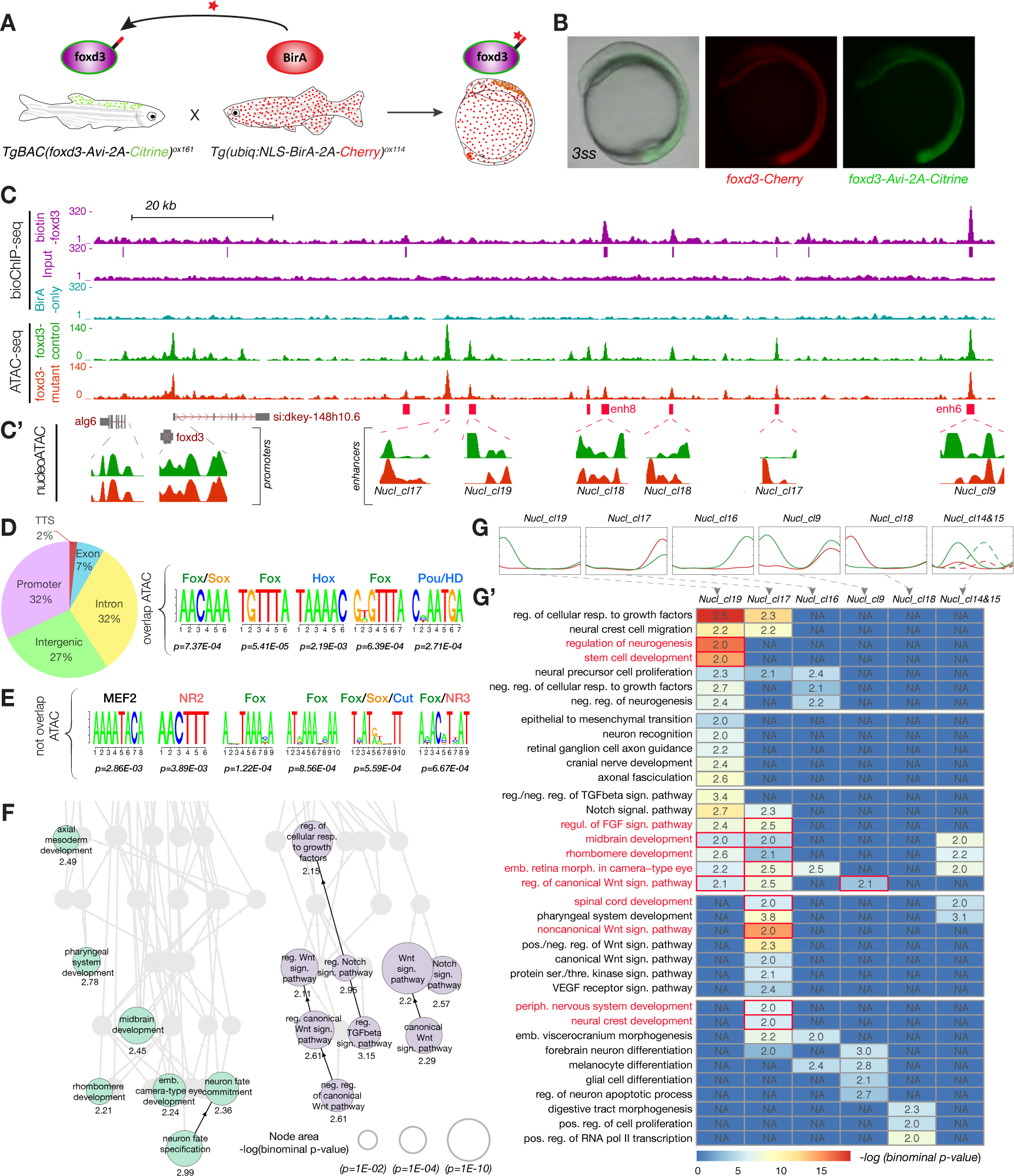
Biotin-ChIP and nucleosomal occupancy analysis. (A) Experimental strategy for biotagging foxd3 protein in vivo. Zebrafish transgenics expressing Avi-tagged foxd3 and ubiquitous NLS-BirA are crossed to obtain embryos expressing biotinylated foxd3 for use in biotinChIP-seq. (B) Lateral view of the embryo issued from crosses of *TgBAC(foxd3-Avi-2A-Citrine)^ox161^* and *Gt(foxd3-mCherry)^ct110R^* shows overlap of Citrine and Cherry reporters. (C) Genome browser screenshot showing mapped biotin-foxd3 ChIP-seq and ATAC-seq within the foxd3 regulatory locus at 5-6ss. BirA-only ChIP-seq control is shown in cyan. Positions of called peaks are indicated as magenta vertical lines underneath biotin-foxd3 ChIP track. (C) Nucleosomal occupancy tracks (NucleoATAC) showing nucleosome positions within the regulatory elements in foxd3 mutant (red) and control (green) cells. (D) Pie chart detailing genomic feature annotations of foxd3-bound regions. Top position weight matrices (PWMs) and associated TFs enriched within Foxd3-bound elements overlapping (D) and not overlapping (E) ATAC-seq peaks. (F) Directed acyclic graph showing hierarchical view of enriched biological process ontology terms for foxd3-bound elements indicates that foxd3 directly controls two independent regulatory aspects – wnt (and notch) signalling and NC neural derivative fates. (G) Merged mean profiles of nucleosomal clusters obtained by *k*-means analysis showing different nucleosomal patterns between foxd3-mutant (CC, red) and control (C, green) neural crest. (G’) Heatmap combining functional annotation by GREAT of elements belonging to different nucleosomal clusters detected at 5-6ss. Ontology terms enriched only on foxd3-bound elements are labelled and boxed in red. See figure S6 for nucleosomal clustering at epiboly.

Comprehensive binding motif maps of foxd3-bound putative enhancers from different clusters suggested that during active NC specification, foxd3 likely co-operates with some core NC factors, such as snail and nr2f (Fig. S6A). However, the enhancers bound by foxd3 at this stage are no longer enriched in soxE, msx or prdm motifs and do not coincide with CTCF. In contrast, these elements are highly enriched in sox2, klf and POU factor binding sites, suggesting that, at later stages, foxd3 is likely directly involved in regulation of the stem cells properties in the migrating/differentiating NC cells.

Altogether, our foxd3 biotinChIP-seq data in premigratory NC argue for a direct activation of the majority of the identified hotspot NC enhancers, particularly those controlling signalling pathways, such as Notch and Wnt. Notably, a subset of foxd3-bound peaks do not overlap with open chromatin regions at 5-6ss, supporting our hypothesis that foxd3 acts as a pioneer factor during NC specification as well as migration.

### foxd3 controls nucleosomal rearrangements at NC elements

Functional annotation of foxd3-bound ATAC-seq positive elements and hierarchical analysis of gene on-tologies using GREAT tool suggested that foxd3 intervenes at multiple regulatory levels in premigratory NC (5-6ss). On one hand, it acts upstream of regulatory factors and components of major signalling pathways (Wnt, Notch and TGF beta), while in parallel controlling fate specification of future NC neuronal and mesenchymal derivatives and preventing other developmental fates (Fig. 6F). The majority of these directly regulated signalling molecules are silenced by the time of active NC migration/differentiation and upregulated in *foxd3*-mutant NC, suggesting a switch to a later repressive role for foxd3. Given that foxd3-bound NC regulatory elements were previously opened, acetylated and likely used during NC specification, we investigated whether foxd3 acted via a supplementary mechanism, via reorganisation of local chromatin in order to rapidly switch between priming and repression. To this end, we generated nucleosomal occupancy tracks (Fig. 6C′) using NucleoATAC algorithm that enables calling nucleosome positions using Tn5 footprints embedded in ATAC-Seq data (Schep et al., 2015). *K*-means clustering identified cohesive groups of elements that presented significant differences in nucleosomal patterns between *foxd3*-mutant (CC) and control (C) NC. Interestingly, while no changes in chromatin architecture at promoters were observed, nucleosomal clustering at 5-6ss singled out groups with differential nucleosomal density in *foxd3* mutants (Fig. 6G). Functional annotation of nucleosomal clusters containing elements that were more compactly organised in *foxd3*-control cells and widely nucleosome-free in *foxd3*-mutants (nucl cl19, 16 and 9) indicated their involvement in repressing major signalling pathways (Notch, FGF, TGF beta), as well as controlling the stem cell programme and preventing premature differentiation of derivatives (neurogenesis) within the peripheral nervous system and the brain (Fig. 6G’). Analysis of elements from these foxd3 bound clusters at 5-6ss indicated that direct foxd3 activity is required for controlling stem cell and neurogenesis programmes in migrating neural crest, as well as to repress regulators of canonical Wnt pathway. This further highlights the importance of temporal dynamics of foxd3 activity and provides insight into which foxd3-regulated functions initiate at the 5-6ss. Clusters with inverted patterns such as nucl cluster 17 containing elements with higher nucleosome occupancy in *foxd3*-mutants yield significant statistical association to actively maintained NC genes as well as to molecules involved in NC migration and formation of peripheral nervous system derivatives that are being primed at this stage. This suggests that foxd3-dependent nucleosomal rearrangements at enhancers also mediate gene activation necessary for the next steps of NC ontogeny. Interestingly, foxd3 appears to positively regulate a different complement of signalling pathway genes (FGF, BMP, VEGF), as well as directly control regulators of non-canonical Wnt pathway genes, necessary for migration via nucl cluster 17 (Fig. 6G’).

Similar analysis of the nucleosome occupancy at the epiboly stage has identified both clusters of elements with activating patterns, which in addition to controlling onset of core NC genes also regulate Notch, FGF and canonical Wnt pathways, as well as clusters with repressive patterns, which act to delay of stem cell differentiation and NC migration programmes (Fig. S6B-D). This shows that the role of foxd3 in priming NC specification is also mediated by nucleosomal re-shuffling.

Foxd3-mediated dynamic re-shuffling of nucleosomes, allows a switch from activation to repression state, thus explaining how some of the core NC genes are first downregulated in *foxd3*-mutants and later overexpressed. We identified more than 1200 elements that are initially nucleosomally primed and later reshuffled to a repressive state. Strikingly, these elements significantly associate with genes upregulated at 14ss (*p*=0.01187999) as well as with primed NC specification genes downregulated at 5-6ss (*p*=0.001845046) and mainly control processes of stem cell and NC development and Delta Notch signalling pathway. Clustering based on the differential metrics (chromatin accessibility, H3K27Ac or nucleosomal occupancy) between *foxd3*-mutant and control cells identified cohesive groups of non-promoter regulatory elements, reflecting multiple roles of foxd3 during NC development. We found that these foxd3 roles, as well as its mechanistic modes of action, are uncoupled, with no clear association between different types of clustered elements (Fig. S7). In particular, the majority of differential accessibility and nucleosome clusters did not correlate with differential H3K27Ac elements, suggesting that H3K27 acetylation was uncoupled from both chromatin opening and nucleosome re-shuffling mechanisms.

On the whole, these results show that foxd3 coordinates nucleosome occupancy primarily at *cis*-regulatory rather than promoter regions. Prior to NC specification, foxd3-mediated reorganisation of nucleosomes helps:(1) prime the elements used for onset of NC specification genes by ensuring their permissive conformation (open in *foxd3*-controls and closed in *foxd3*-mutants) and (2) repress/close elements of NC genes only expressed at later stages. Similarly, at 5-6ss, foxd3 opens enhancers that promote stem cell maintenance, the migration apparatus and appropriate NC cell fates. Concomitantly, foxd3 represses unwanted cell fates, stem cell differentiation and key signaling pathways like the canonical Wnt pathway. Nucleosome re-shuffling is likely to be the major mechanism by which foxd3 exercises its bimodal activity, switching from a pioneer to a repressor depending on the developmental context and stage of NC development.

### Increasing resolution of the neural crest gene regulatory network

We used Weighted Gene Co-expression Network Analysis (WGCNA) (Langfelder and Horvath, 2008) of *foxd3*-expressing and *foxd3*-negative datasets to identify clusters of genes with highly correlated expression dynamics (Fig. 7A). After generating signed weighted gene networks (Mason et al., 2009), we analysed the co-expression modules corresponding to genes positively and negatively correlating with *foxd3*+ cells (Fig. 7A-B). The results show that identified clusters largely retraced the structure of the previously proposed NC-GRN.

**Figure 7.**
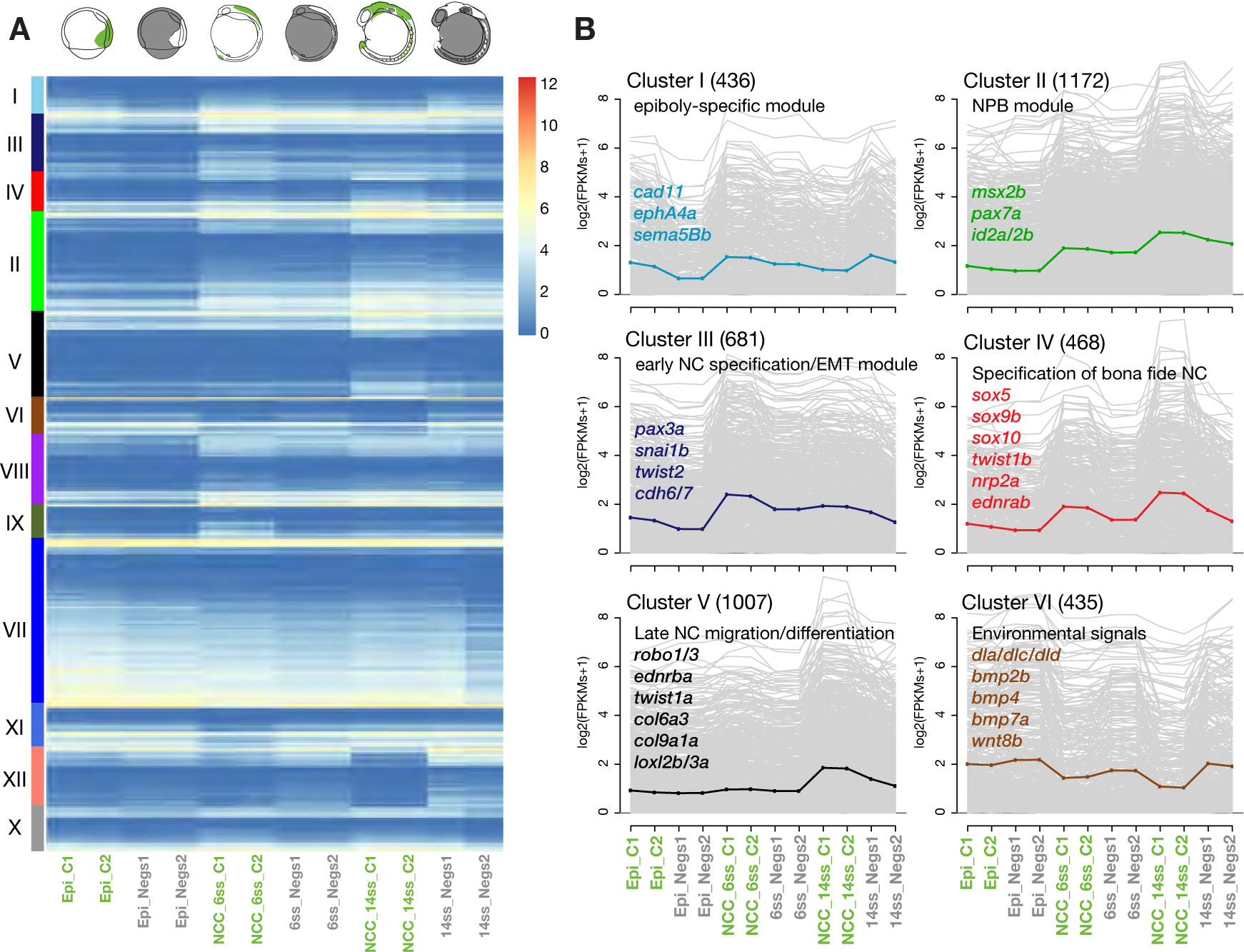
(A) Heatmap of expression profiles across *foxd3*-expressing and *foxd3*-negative datasets at 75% epiboly, 5-6ss and 14ss for genes in 12 modules within the obtained signed network. (B) Plots depicting log-transformed expressions levels of genes consistently co-expressed in *foxd3*-expressing or non-expressing cells across developmental time and organised into top assigned modules (Clusters I-XII). Clusters retrace the previously proposed modular structure of NC-GRN. Average expression level curves representing module pattern are shown in the same colour as gene symbols for NC genes previously assigned to corresponding NC-RNGRN modules. (See Fig. S7 and ShinyApp for membership assignment for all modules).

The main exceptions from previous network description were in Cluster I that consisted of genes highly expressed and enriched at 75% epiboly and in 5-6ss premigratory, but not migrating NC at 14-16ss. Although statistical overrepresentation analysis yielded gene ontology (GO) terms reflecting biological processes of signal transduction and cell communication in Cluster I, this novel module did not comprise signalling factors responsible for the induction at the neural plate border, as proposed by the NC-GRN model (Meulemans and Bronner-Fraser, 2004), but included signals and cell surface receptors important at later stages of NC ontogeny (*cadherin 11 type 2, eph receptor A4a*, neural adhesion molecule, *nadl1.2, semaphorin 5Bb*, etc.). Hence, we did not recover a single cluster depicting the induction process at the border. Rather, signalling molecules involved in initial steps of neural crest formation were distributed across several clusters, with Notch homologues, as well as notch-regulated proteins that positively correlated with *foxd3*+ cells found in Cluster VII (*notch2, notch3, stno2a, nrarpb*).

Conversely, canonical Notch ligands, deltas, (*dla, dlc* and *dld* in Cluster VI and *dll4* in Cluster XII), as well as bmp homologues (*bmp2b, bmp4* and *bmp7a* in Cluster VI) were recovered as a part of negatively correlating clusters, suggesting they act as environmental cues, emanating from surrounding tissues. Cluster II, characterised by sharp increase in gene expression over the developmental time, largely recapitulated the previously postulated neural plate border (NPB) module, comprised of specification genes (*msx2b, pax7a, id2a/2b*, etc.). It also included recently reported NC regulators *nr2f2/nr2f5* (Rada-Iglesias et al., 2012), a previously characterised regulatory circuit (*hoxa2, hoxb3, hoxa3, hoxb3, epha4b* and *mafba*) that controls segmentation, patterning and migration of hindbrain NC (Trainor and Krumlauf, 2000), and precise signalling components co-regulated with border specifiers (*wnt6a, sfrp2, fzd2, fgf2, fgf19, notch4*). The NC specification module was covered by 2 WGCNA clusters: Cluster III that contained early neural crest specifier genes, such as *pax3a* and *snai1b*, but also many molecular components essential for epithelial to mesenchymal transition in the NC (*cad6/cad7, twist2, olfml2a* and *olfm3b*) and Cluster IV, containing *bona fide* NC specifiers (*sox5, sox8a, sox9b, sox10, twist1b, zeb2a* etc.), characterised by the strong specific signal in NC starting from 5-6ss. This module also comprised cell surface receptors, such as *nrp2a, ednrab* and *pdgfra*, essential for neural crest emigration and specification. Factors enriched in NC specifically at the later stage (14ss *foxd3*+ cells) that reflect the role in active migration (*robo1, robo3, ednrba, ets1*), but also involvement in differentiation into specific derivatives (*twist1a, col9a1a, col6a3* and lysyl oxidase-like factors, *loxl2b* and *loxl3a*) were contained in Cluster V, which was equivalent to the NC differentiation module in the NC-GRN. For detailed cluster analysis and description see Fig. S7B. The full content of clusters and regulatory co-expression modules can be explored using the ShinyApp.

## Discussion

Gene expression is the product of interplay between proximal and distal *cis*-regulatory elements, controlling competence at the chromatin level (Ong and Corces, 2012; Wang et al., 2015). Moreover, broad epigenetic changes to the *cis*-regulatory landscape, including histone and DNA demethylation, histone acetylation and loss of heterochromatin characterise different stages of transition from näive to primed pluripotency (Krishnakumar and Blelloch, 2013). Several mechanisms explaining how Foxd3 promotes pluripotency *in vitro* have been proposed. Foxd3 can recruit Tle4, to repress differentiation-associated genes induced by NFAT signalling, through regulation of histone de-acetylation (Zhu et al., 2014). In two recent studies investigating the transition from ESCs to EpiCs, EpiLCs and PGCKs, mouse Foxd3 was implicated in the regulation of stem cell pluripotency by associating to different enhancer marks and subsequently manipulating transcriptional competency of downstream genes (Liber et al., 2010). The first report showed that Foxd3-bound enhancers associated with genes primed for expression upon exit from näive pluripotency, with Foxd3 promoting nucleosome depletion by recruiting SWI/SNF complex chromatin remodeller Brg1, whilst simultaneously acting as a repressor and preventing enhancer acetylation by recruiting HDACs (Krishnakumar et al., 2016). The other study showed that Foxd3-bound active enhancers associated with highly expressed genes that become silenced upon exit from näive pluripotency, where corresponding enhancers were decommissioned through recruitment of Lsd1, and a reduction in p300 activity (Respuela et al., 2016). Surprisingly, the two studies found a minimal overlap (only 12%) in FoxD3 bound peaks (Plank et al., 2014; Sweet, 2016; Yong et al., 2016). The discrepancies between the different putative mechanisms of FoxD3 re-enforced the need for *in vivo* studies that would characterise the regulatory context within which FoxD3 mediates different activator and repressor roles across developmental time.

### Developmental functions of FoxD3 in the neural crest

In vertebrate embryos *foxd3* is expressed in premigratory NC, maintained in early migrating NC and downregulated in cranial and cardiac NC derivatives except for glial lineages (Dottori et al., 2001; Kos et al., 2001; Mundell and Labosky, 2011; Nitzan et al., 2013b). Previous *foxd3* knockdown studies in zebrafish mainly described the phenotypes affecting NC derivatives, reporting defects in the specification of cartilage progenitors and pigment cell precursors, as well as delay in differentiation of pigment and neural progenitors (Lister et al., 2006; Montero-Balaguer et al., 2006; Stewart et al., 2006). Similarly, *foxd3* knockout using foxd3 Cherry mutant homozygotes showed persisting mesenchymal phenotype at post-migratory stages and an increase in melanocytes in the trunk (Hochgreb-Hägele and Bronner, 2013). The co-operative activity of foxd3 with another NC specifier, tfap2a, was shown to be critical for NC induction, with two factors acting in concert to fine-tune Wnt and Bmp signalling gradients during gastrulation (Wang et al., 2011). Interestingly, in the chicken embryo, Foxd3 is required for expression of NC specifier genes, but was also independently involved in controlling EMT by downregulating the transmembrane scaffolding protein Tetraspanin18, prior to NC emigration (Fairchild et al., 2014). Finally, FoxD3 has been highlighted as one of the central factors in maintaining NC cell stemness, but precise regulatory mechanisms mediating this role have not been elucidated (Mundell and Labosky, 2011).

### FoxD3 is a Pioneering factor for NC specification

These studies suggest foxd3 plays an array of complex independent roles during NC ontogeny, but its role during NC specification has remained elusive. Although Foxd3 was thought to act mostly as a transcriptional repressor, previous studies failed to recover more differentially upregulated versus downregulated genes in *foxd3* mutant cells (Respuela et al., 2016; Yaklichkin et al., 2007). Strikingly, our analysis showed that foxd3 plays a central activating role in NC specification, controlling the expression of an entire NC specification module. We present evidence that foxd3 acts at a global level to prime NC factors by modulating the accessibility of their *cis*-regulatory elements. Thus, much like its relatives, FoxA1 and FoxA2, shown to regulate enhancer dynamics for specific gene expression controlling pluripotent stem cell potential, cell fate transitions, lineage choice and differentiation (Adam et al., 2015; Serandour et al., 2011; Zaret and Carroll, 2011), foxd3 acts as a pioneer factor in the neural crest. By studying dynamics of chromatin opening across several stages, we identified a set of hotspot enhancers whose accessibility was dependent on foxd3. Quantification of accessibility levels using normalised ATAC assay and statistical differential binding analysis indicated that defects in *foxd3*-mutant cells are most striking at the onset of enhancer opening and affected early genes at the onset of NC specification, late genes at the onset of migration and genes involved in the maintenance of stemness.

### Foxd3 affects H3K27 Acetylation on NC enhancers

Previous studies suggested that one of the *modi operandi* of pioneer factors was the recruitment of H3K27 acetyltransferase activity, a hallmark histone modification of active enhancers (Choi et al., 2016; Kerschner et al., 2014). In contrast, a recent report found that following FoxA1/A2 activity, accessible nucleosomes in liver-specific enhancers had reduced H3K27Ac, suggesting that the initial role of pioneer factors in opening and controlling nucleosome occupancy at enhancers was temporally uncoupled from the acetylation role (Iwafuchi-Doi et al., 2016). We found that lack of *foxd3* during NC specification resulted in differential K27 Acetylation, with some NC regulatory elements showing depletion and others an increase in H3K27Ac mark in mutant embryos. We show that early NC specifiers, downregulated in *foxd3*-mutants, are controlled positively via this mechanism, as they associated to the K27Ac-depleted elements with a high statistical significance. At the same time, we demonstrate that those *cis*-regulatory elements, which show significant areas of hyperacetylation in mutants, negatively control essential components of Wnt signalling pathway. Thus, foxd3 may act to decommission enhancers and hence repress factors that need to be downregulated for NC migration/differentiation to proceed.

### Foxd3 affects nucleosomal positioning on NC enhancers

FoxD3 forkhead DNA binding domain, like that of FoxA proteins, is constituted of three helices and two large loops (wings) and is remarkably similar to the winged-helix structures of linker histone H1 that avidly binds nucleosomes (Clark et al., 1993). Such pioneer factors have been suggested to induce nucleosome repositioning, possibly by recruiting hyperdynamic histone variants, like H2A.Z and H3.3 and other chromatin and DNA modifying proteins, to allow binding of *cis*-regulatory elements by transcriptional complexes (Chen and Dent, 2014; Spitz and Furlong, 2012; Zaret and Carroll, 2011). Our analysis shows that although it does not affect nucleosomal occupancy at promoters, foxd3 influences the nucleosome positioning of NC regulatory elements in a context-dependent manner, resulting in both ‘permissive’ and ‘repressive’ chromatin organisations. Clustering of these elements at different stages, based on differential nucleosomal patterns, shows that this foxd3-controlled mechanism is both used to prime NC specification, migration/differentiation and stem cell genes, as well as to prevent premature differentiation and directly repress components of signalling pathways no longer active in migrating crest.

### Biotin-ChIP confirms direct binding of foxd3 to NC regulatory elements

Analysis of elements directly bound by foxd3 demonstrates that one of the primary functions of foxd3 in premigratory NC is to directly control components of essential signalling pathways, Notch, TGF beta and in particular Wnt. This role is highlighted throughout different stages of our analysis, as crucial signalling molecules are downregulated as NC development progresses, are upregulated (de-repressed) in foxd3 mutants, and their regulatory elements are bound by foxd3 and decommissioned. At this stage, foxd3 also directly positively controls factors involved in differentiation of neuronal/peripheral nervous system derivatives and components of non-canonical Wnt signals, essential for NC migration. Last, but not least, foxd3 binds to regulatory elements controlling the stem cell programme by both directly priming genes involved in the maintenance of stem cell potential and directly repressing genes involved in stem cell differentiation, thus overall maintaining NC multipotency.

### Bimodal action of foxd3

Here we present striking evidence that during neural crest formation *in vivo*, in addition to its canonical role as a repressor (Yaklichkin et al., 2007), foxd3 acts as a pioneer factor to prime NC gene expression. In line with recent *in vitro* studies (Krishnakumar et al., 2016; Respuela et al., 2016), we demonstrate that foxd3 functions primarily by changing the chromatin landscape of *cis*-regulatory elements and that its different modes of action are functionally uncoupled. We show that, in the developing embryos, foxd3 is capable of pioneering NC chromatin regulatory landscape in a bimodal fashion, facilitating both permissive and repressive states. These mechanisms do not exhibit sharp temporal boundaries, but instead, they are present simultaneously, with a gradual shift towards the repressive activity after NC specification. Such bimodal activity of foxd3 enables early NC fate transition and maintenance of multipotency. Overall, we show that foxd3 sets up the core NC gene enhancers (Cluster 3.1), as well as later migratory NC regulatory elements required for the specification of distinct NC lineages. As a chromatin silencer, it negatively regulates mesenchymal gene expression preventing pre-mature NC differentiation and shuts down signalling pathways that need to be downregulated to enable NC migration and differentiation. We present striking evidence that during NC ontogeny foxd3 may switch from permissive to repressive nucleosome/chromatin organisation using same regulatory elements. At the same time, we also show that the bimodal foxd3 activity is involved in fine-tuning the gene expression by employing different regulatory elements for the same gene (from Clusters 3.1 and 3.2 for instance) to mediate activating versus repressing regulatory activity. The net result of foxd3-associated chromatin state is likely to be dependent on its *in vivo* interacting partners, such as other TFs or chromatin remodellers, which need to be investigated in the future.

### Elaboration of the NC-GRN

The wealth of genome-wide regulatory data generated in this study offers an exquisite opportunity to expand and build a highly detailed representation of proposed NC gene regulatory network (NC-GRN). We exploited our NC-specific bulk transcriptome data using WGCNA to extend current GRN modules, mainly based on candidate gene approaches and studies of individual enhancers (Barembaum and Bronner, 2013; Betancur et al., 2010b; Simoes-Costa et al., 2012). WGCNA clustering revealed modules of co-regulated genes that closely reflected previously established NC-GRN structure, particularly at the level of the specification of *bona fide* NC. The full complement of factors and their position within the NC-GRN modules can be surveyed using ShinyApp. Although our transcriptome data, obtained from a small number of cells collected from clutches of sibling embryos expressing fluorescent NC marker, does represent NC gene expression at precise time points, it also incorporates both temporal and biological variation within the single embryo and across the population. As such, it cannot be used to ascertain the co-presence of studied NC genes with their upstream regulatory factors. We, therefore, used our scRNA-seq data which has exquisite resolution and features high coverage/sequencing depth and minimal dropout rate, to generate the single cell expression catalogues to screen for genes co-expressed in the same cell and thus participating in individual gene regulatory circuits (gene expression cut-off RPKM*>*1). Finally, we present an integrated, user-friendly ShinyApp enabling access and use of all the regulatory information obtained in this study. This application facilitates interrogation of gene regulatory interactions within the foxd3-primed minimal core NC specification module, by connecting a full complement of associated *cis*-regulatory elements, corresponding enriched TF binding maps, tightly controlled NC specification target genes and their co-expression modules. Thus our study provides a platform for building holistic representations of NC-GRN kernels using genome-wide regulatory data and expanding our understanding of the NC-GRN.

## Author contributions

Conceptualisation, T.S.S., D.G.; Methodology, R.M.W., M.L., V.C.M.; Software, D.G., E.R.; Validation, T.S.S., D.G., R.M.W., M.L.; Formal Analysis, D.G., T.S.S., S.T.; Investigation, R.M.W., M.L., V.C.M., U.S., T.H.H.; Writing - Original Draft, T.S.S.; Writing Review & Editing, T.S.S., D.G., R.M.W., M.L., V.C., S.T.; Visualisation T.S.S., D.G.; Supervision, T.S.S.; Funding Acquisition, T.S.S. and A.M.

## Acknowledgements

This work was supported by MRC (G0902418), Lister Institute prize, Leverhulme Trust grant (RPG-2015-026), March of Dimes Basil OConnor Award to TSS, and SNF Fellowship to DG.

## Experimental Procedures

### Zebrafish Husbandry

Animals were handled in accordance to procedures authorized by the UK Home Office in accordance with UK law (Animals [Scientific Procedures] Act 1986) and the recommendations in the Guide for the Care and Use of Laboratory Animals. All vertebrate animal work was performed at the facilities of Oxford University Biomedical Services. Adult fish were maintained as described previously (Westerfield, 2000). Embryos were staged as described previously (Kimmel et al., 1995).

### Cell dissociation and FAC-sorting

Embryos were sorted by fluorescence, pooled and dissociated with collagenase (20mg/ml in 0.05% trypsin) to a single cell suspension, centrifuged and re-suspended in Hanks buffer. Fluorescent positive cells were sorted and collected using BD FACS-Aria Fusion.

### Bulk RNA extraction, library preparation and sequencing

FACS sorted cells were washed with PBS and stored at −80°C in lysis bu er. RNA was extracted using Ambion RNAqueous Micro Total RNA isolation kit (AM1931), checked on Bioanalyser, samples with RIN *>*7 were used to prepare cDNA using Takara Clontech SmartSeq2 V4 kit (634889). Sequencing libraries were prepared using Illumina Nextera XT library preparation kit (FC-131-1024). 75% Epiboly-stage cell libraries (Citrine-expressing, Citrine-Cherry-expressing and cells not expressing *FoxD3*) were sequenced using 80 bp reads using Illumina Nextseq500 platform. 5-6ss cell libraries expressing *FoxD3* (Citrine-expressing, Citrine-Cherry-expressing) were sequenced using 50bp reads using Illumina Hiseq2000 platform, and cells not expressing FoxD3 using 80bp reads on Illumina Nextseq500 platform. 12ss cell libraries expressing *FoxD3* (Citrine-expressing, Citrine-Cherry-expressing) were sequenced using 100bp reads using Illumina Hiseq2000 platform. 14ss cell libraries (Citrine-expressing, Citrine-Cherry-expressing and cells not expressing FoxD3) were sequenced using 80bp reads using Illumina NextSeq500 platform. All sequenced libraries were paired end.

### H3K27Ac ChIP, library preparation and sequencing

FACS sorted cells were cross-linked with 1% formaldehyde and sonicated to 300-800bp fragments. Pre-blocked Protein A Dynabeads were incubated with antibody (Abcam Ab4729). Sonicated DNA-protein complexes were applied to beads, IgG antibody (Millipore 12-370) was used as control and an input sample was taken. Samples were washed, cross-links were reversed, immunoprecipitated DNA was eluted, purified and libraries prepared using amplification protocol for small cell numbers (Adli and Bernstein, 2011). H3K27Ac ChIP libraries were sequences using 50bp paired end reads using Illumina Hiseq2500 platform.

### Generation of Avi-tagged foxd3 transgenic line

Tol2-mediated BAC-mediated transgenesis, as described in (Trinh et al., 2017), was used to generate *TgBAC(foxd3-Avi-2A-Citrine)^ox161^* transgenic line. Avi-2A-Citrine recombination donor construct was generated by PCR using Pfu polymerase (Pfu UltraII Hoststart PCR Master Mix, Agilent Technologies) and cloned using InFusion (InFusion HD Cloning kit, Clontech). It contains a FLAG epitope, a TEV protease recognition sequence, an in-frame 48 bp long Avi-Tag and a fluorescent reporter Citrine followed by a polyA tail with a kanamycin selection cassette flanked by flippase recognition target (FRT) sequences. Citrine reporter is separated from the foxd3-Avi gene by a linker viral 2A sequence that mediates a ribosome skipping, thus allowing for co-expression of both components from a single transcript (Kim et al., 2011). Genomic context of the *Danio rerio* BAC clone CH211-196F13 (203kbp) was used for recombineering, as it harbours a full single exon ORF of the *foxd3* gene and the upstream DNA regions of more than 200kbp, allowing to capture not only the *foxd3* promoter region but also the associated *cis*-regulatory elements. A recombination cassette (foxd3-Avi-2A-Citrine) containing 5′ and 3′ homology arms to the *foxd3* locus was used to replace a wild-type *foxd3* gene with the Avi-tagged version. NLS-BirA zebrafish transgenic line *Tg(ubiq:NLS-BirA-2A-Cherry)^ox114^* expresses 3 x HA epitope, nuclear localisation signal (NLS) sequence, BirA gene sequence, viral 2A sequence and a Citrine reporter gene under the control of ubiquitous ubb promoter.

### FoxD3 Biotin-ChIP, library preparation and sequencing

Foxd3 Biotin-ChIP was performed on 320 5-6ss whole embryos (128,000 cells of interest). Embryos were manually dechorionated, cells were dissociated with 20 strokes using pestle A in isotonic nuclei extraction buffer (NEB: 0.5% NP40, 0.25% Triton X, 10 mM Tris-HCl (pH 7.5), 3 mM CaCl2, 0.25 M sucrose, 1mM DTT, 0.2 mM PMSF, 1X Proteinase inhibitor (PI)) in a glass homogeniser and cross-linked using 1% formaldehyde at room temperature for 10 min. Fixation was quenched with 125 mM of glycine for 5 min. Cross-linker was washed-out by 3x pellet washes with 1x PBS (with 1X PI, 1 mM DTT and 0.2 mM PMSF) centrifuging at 2000 × g for 4 min at 4°C. Pellets were re-suspended in NEB. Cell nuclei were expulsed with 20 strokes using pestle B in a glass homogeniser, pelleted and washed with 1 xPBS (with 1X PI, 1 mM DTT and 0.2 mM PMSF). Nuclei were lysed in SDS lysis buffer (0.7% SDS, 10mM EDTA, 50 mM Tris-HCl (pH 7.5), 1x PI). Cross-linked chromatin was sonicated at 12A 10x (10s ON 30s OFF) followed by 8A 4x (30s ON 30s OFF). Sheared chromatin samples were pre-cleared in pre-blocked Protein G beads (Dynabeads Protein G, Life Technologies) for 1 hour at 4°C. 1/20 of biotinChIP was collected as an input fraction and stored at −80°C. Pre-cleared chromatin samples were incubated on pre-blocked streptavidin beads (Dynabeads M-280 streptavidin beads, Life Technologies) o/n at 4°C. Beads were washed with SDS Wash Buffer (2% SDS, 10mM Tris-HCl (pH 7.5), 1 mM EDTA) at room temperature, followed by 4x RIPA washes (50 mM Hepes-KOH (pH 8.0), 500 mM LiCl, 1mM EDTA, 1% NP40, 0.7% Na-Deoxycholate, 1x PI) and 1x NaCl TE wash (1x TE, 50mM NaCl) at 4°C. Chromatin was eluted from the beads with SDS ChIP elution buffer (50 mM Tris-HCl (pH 7.5), 10 mM EDTA, 1% SDS). Cross-linking was reversed o/n at 70°C in the thermomixer at 1300 rpm. Cellular RNA was digested with RNAse A (0.2 g/ml) at 37°C for 1 hour, and cellular protein with Proteinase K (0.4 mg/ml) at 55°C for 2 hours. Chromatin samples were separated from the streptavidin beads. Input and ChIP DNA was extracted using a standard phenol-chloroform extraction method. Libraries were prepared using MicropPlex Library Preparation kit (Diagenode) and sequenced using NextSeq^®^ 500/550 High Output Kit v2 (75 cycles) on NextSeq 500 sequencing system.

### ATAC, library preparation and sequencing

FACS sorted cells were lysed (10mM Tris-HCl, pH7.4, 10mM NaCl, 3mM MgCl2, 0.1% Igepal) and tagmented using Nextera DNA kit (Illumina FC-121-1030) for 30 minutes at 37 °C. Tagmented DNA was amplified using NEB Next High-Fidelity 2X PCR Master Mix for 11 cycles. Tagmentation efficiency was assessed using Agilent Tapestation. ATAC-Rx was carried as previously described (Orlando et al., 2014), and as described above with the addition of 50% extra *Drosophila* S2 cells. ATAC-seq libraries were sequenced using paired-end 40 bp run using Illumina NextSeq^®^ 500 platform.

### Single cell RNA preparation, library preparation and sequencing

Individual cells were collected by FACS, cDNA was obtained and sequencing libraries prepared as previously described (Picelli et al., 2014). Libraries were sequenced using 50 bp single end reads for 96 cells. 4x10^7^ dilution of ERCC spike in control was used.

### Enhancer reporter constructs

All enhancer inserts were generated by PCR using KAPA Long Range HotStart PCR kit (Kapa Biosystems) and cloned into the E1b:GFP:AC-DS vector using the InFusion kit (InFusion HD Cloning kit, Clontech). Fertilised single-cell embryos were injected with 30 pg of plasmid DNA and 25 pg of Ac mRNA. Injected embryos were imaged on a Zeiss780 LSM inverted confocal microscope equipped with EC Plan-Neofluar 10x/0.30 NA WD=5.2 (Zeiss) objective or using a Zeiss Axio Scope.A1 equipped with 5x/0.15 NA N-Achroplan or 10x/0.3 NA EC Plan-Neofluar objectives (Zeiss) at desired developmental stages.

### In situ hybridisation

*In situ* hybridisation was performed with anti-sense RNA probes according to standard protocols, as described previously (Hochgreb-Hägele and Bronner, 2013). Probe synthesis was conducted with the components of the DIG RNA Labelling Kit (Roche).

### Bioinformatic Processing

#### Bulk RNA-seq Processing

Reads were trimmed to remove low quality bases using sickle (Joshi and Fass, 2011) when necessary. Read quality was evaluated using FastQC (Barbaraham). Mapping to GRCz10/danRer10 assembly of the zebrafish genome downloaded from UCSC Genome Browser was performed using STAR 2.4.2a.(2) (Dobin et al., 2013). Read counts were obtained using subread FeatureCounts (v1.4.6-p4) (Liao et al., 2014) using standard parameters using a gene model gtf derived from Ensembl annotation downloaded from UCSC genome browser. Gene model for ENSDARG00000095311 (the antisense transcript of FoxD3), was removed from gene models. Differential Expression analysis was carried out using in DESeq2(v.1.14.1).

### Analysis of single-cell RNA sequencing

Short reads (51bp) from 96 cells were aligned to the zebrafish genome (GRCz10/danRer10 assembly) and ERCC spike-in controls using STAR (Dobin et al., 2013) with default parameters. The featureCounts (Liao et al., 2014) was then used to count the number of mapped reads to the reference gene models. Expression values were quantified as read per kilobase of transcript length per million of mapped reads (RPKM) on the basis of Ensembl gene annotation using the rpkm function in edgeR (Robinson et al., 2010). We used cells with higher than 100,000 mapping reads and 2,000 detected genes (RPKM *>*1) for the downstream analysis. With these cut-off criteria, one cell was excluded due to the low sequencing depth. We performed the principal component analysis (PCA) using the custom R script. We selected top 500 genes with the highest absolute correlation coefficient (PCA component loadings) in one of the first three components and then performed PCA and T-distributed stochastic neighbour embedding (tSNE) analyses. The heatmap was visualised on selected gene sets based on the log2 of RPKM scale using the heatmap function in R. For purpose of single cell transcriptional cataloguing, the scRNA-seq data is visualised using SCDE package (http://hms-dbmi.github.io/scde/index.html) (Fan et al., 2016). Additional analysis was carried out using PAGODA R package (Fan et al., 2016). 50% epiboly demultiplexed sc-RNASeq data was kindly provided by R.Satija (Satija et al., 2015), and processed as described above.

### Differential expression at 50% epiboly

Robustness of differential expression results in these cells using a resampling approach similar to bootstrap: expression values were iteratively randomly pooled in samples of 100 FECs and 100 FNECs and used for differential expression analysis (1000 iterations). By counting the number of times each gene was found differentially enriched or depleted (bootstrap value), we identified genes that were differentially expressed in FECs (fdrtools, adjusted P-value¡0.05) in a manner comparable to the analysis carried out on bulk RNA-seq samples ATAC-seq Processing Reads were trimmed for quality using sickle when necessary and mapped using bowtie (v.1.0.0) (Langmead et al., 2009). Bigwig files were generated using an enhanced Perl script courtesy of Jim Hughes. Peak calling was performed as described previously (Buenrostro et al., 2013). Briefly, bam files were sorted by name and paired end bed files were obtained using bedtools(v.2.15.0) bamtobed bedpe. Reads that were not properly paired were discarded and paired reads were displaced by +4 bp and −5 bp. Reads were extended to a read length of 100bp.Peak calling was performed using MACS2 callpeak f BED shiftsize=100 nomodel slocal 1000 parameters (Zhang et al., 2008). To obtain mappability data, a synthetic 40bp-long single end fastq dataset was generated and mapped using bowtie(v.1.0.0) using m 1 parameter. Bedgraph files were obtained using bedtools genomeCoverageBed -bg split function. MACS2-called peaks that overlapped with regions which in the mappability did not correspond to read size (40bp) were discarded. Identification of peaks corresponding to TSS/promoter, intergenic, intronic and TES locations was carried out using Homer (v.4.8)(Heinz et al., 2010) annotatePeaks.pl script. Only peaks present in both replicates were retained, using bedtools intersect function to generate reference ATAC-seq ensembles for each stage. ATAC-Rx-seq was processed similarly with the exception that a combined genome of containing danRer10 and Drosophila melanogaster dm6 genomes was created and all reads were mapped to the latter. Zebrafish read counts were normalized as described previously (Orlando et al., 2014). *K*-means clustering of ATAC-seq signal was carried out using SeqMINR software as described (Ye et al., 2011), using non-promoter reference ATAC-seq peaks form 5-6ss samples. Nucleosome localization was carried out using nucleoATAC suite using default parameters in peaks called at each stage. Bedtools was used to generate bigwig files and clustering of nucleoATAC bigwig signal was carried out using deepTools (v.2.2.2) using *k*-means clustering with 20 clusters.

### H3K27Ac-ChIP Processing

Reads were trimmed for quality using sickle when necessary and mapped using bowtie (v.1.0.0). Bigwig files were generated using an enhanced Perl script courtesy of Jim Hughes. MACS2 was used to identify peaks using standard parameters. Only peaks present in both replicates were retained, using bedtools intersect function. *k*-means clustering of H3K27Ac signal was carried out using SeqMINR software as described (Ye et al., 2011).

### FoxD3 Biotin-ChIP Processing

Reads were trimmed for quality using sickle when necessary and mapped using bowtie (v.1.0.0). Peak calling was performed using Homer (v.4.8) (Heinz et al., 2010) findPeaks script using size 150minDist 1000 parameters. Bigwig files were generated using an enhanced Perl script courtesy of Jim Hughes. Motif discovery and characterisation was performed using MEME suite (Bailey et al., 2015). Newly discovered motifs obtained using MEME tool were subsequently analysed by TOMTOM tool to identify statistically significantly similar motifs in known transcription factor binding site motif databases (Gupta et al., 2007).

### Transcription Factor Binding Site Identification

Initial Transcription Factor Binding Site (TFBS) enrichment analysis of known motifs was performed using Homer suite (findMotifsGenome.pl) (Heinz et al., 2010). The analysis was performed for all *k*-means clusters, using the default 200bp window centred on the ATAC-peak, and all non-promoter putative regions were used as background. Due to paucity of available zebrafish transcription factor binding sites (TFBS), a clustering approach of known transcription factors sites was utilized. TFBS for each gene family of interest were downloaded from CIS-BP (http://cisbp.ccbr.utoronto.ca) (Weirauch et al., 2014). Binding sites were clustered using gimme suites cluster option (v. 0.9.0.3) (van Heeringen and Veenstra, 2011).Background values for each of the clustered motifs was obtained using gimme background function. Cutoff values relative to background sequences were obtained using gimme threshold function. Binding sites were identified using gimme scan function using threshold values obtained from previous step in peaks obtained form ATAC-seq processing.

### K-means clustering

*K*-means clustering was performed using the R platform (Ye et al., 2011), by applying the linear enrichment clustering approach to the normalised ATAC-seq datasets and computing the accessibility signal over the non-promoter peaks (+/−1.5 kb from the centre) using the ensemble of peaks containing both elements common all C replicates, as well as elements common to all CC replicates as a reference. Differences in chromatin accessibility for different *k*-means clusters were quantified by plotting the normalised C and CC ATAC-seq counts for all of putative regulatory elements in cluster and calculating Pearson correlation coefficients. *K*-means clustering investigating dynamics of chromatin opening at the NC *cis*-regulatory elements was performed on 75% epiboly and bud stage ATAC-seq datasets, using called 5-6ss non-promoter ATAC peaks as a reference. Functional annotation of each *k*-means cluster was performed using the GREAT Tool (McLean et al., 2010), using whole genome as background. GREAT employs annotations of putative *cis*-regulatory elements to nearby genes and their statistical integration to infer their function. Statistical significance of associated terms was calculated using binomial and hypergeometric tests and either Bonferroni of False Discovery Rate correction.

### Diffbind Differential chromatin accesibility analysis

The differential chromatin accessibility analysis of ATAC-seq dataset in *foxd3*-mutant and control conditions was performed using DiffBind package for differential binding analysis of ChIP-seq (Stark R, 2011). Related plots were generated in R. Significantly differentially accessible peaks were identified using the edgeR package, using a reference ATAC-seq peak ensemble. BenjaminiHochberg multiple testing correction of the resulting p-values was used to derive false discovery rates (FDRs) and only differentially accessible elements with an FDR*<*0.1 were taken in account.

### WGCNA analysis

Weighted correlation network analysis (WGCNA) (Langfelder and Horvath, 2008) was performed on normalised gene count tables generated by DESeq2 (Love et al., 2014), according to the pipeline detailed in the on-line WGCNA tutorial (https://labs.genetics.ucla.edu/horvath/CoexpressionNetwork/Rpackages/WGCNA/).

### ShinyApp and single cell catalogues access

The beta version of the ShinyApp associated with the data produced in this study and Pagoda App (Fan et al., 2016) presenting single cell catalogues can be downloaded from https://github.com/tsslab/foxd3.

